# Nutrient Availability as a Driver of Transcriptome Reprogramming in Osteosarcoma

**DOI:** 10.64898/2026.11.05.764667

**Authors:** Jakub M. Tomczak, Zuzanna Senkowska, Ewelina Węglarz-Tomczak

## Abstract

**Background:** Gene expression in mammalian cells is intricately regulated by environmental and metabolic cues. Among these, nutrient availability represents a key feature of the tumor microenvironment and has recently emerged as a potent modulator of oncogenic signaling networks. In this study, we employed a cell-line model of osteosarcoma, a rare but highly aggressive bone malignancy affecting both humans and dogs. We demonstrate that altering the concentration of a single nutrient can profoundly reprogram gene expression in osteosarcoma cells, revealing a previously underexplored phenomenon: nutrient-dependent gene regulation. This finding highlights potential opportunities for therapeutic intervention and underscores the broader implications of metabolic regulation for personalized medicine and nutrition.

**Methods/Results:** We performed bulk RNA sequencing of an established osteosarcoma cell line cultured under varying concentrations of two key nutrients, glucose and glutamine. Differential gene expression analysis revealed a pronounced nutrient-dependent transcriptional response. The identified genes were functionally associated with pathways involved in cancer, development, nervous system regulation, metabolism, cellular signaling, and immune responses. We identified seven distinct expression patterns, one of which exhibited a striking correlation with cell growth rate and resembled a FOS-like transcriptional signature. Notably, the proto-oncogene *FOS* was among the identified genes and displayed a strong growth-rate-dependent expression pattern.

**Conclusion/Impact:** Our findings demonstrate that modulation of nutrient availability profoundly reshapes the gene expression landscape of osteosarcoma cells, revealing nutrient-responsive targets with potential therapeutic relevance. This work expands our understanding of metabolic regulation in cancer and suggests that nutrient-based interventions may provide a controllable means of influencing tumor progression. Beyond oncology, such approaches may also have broader implications for developmental biology, autoimmune disorders, and nervous system function. More broadly, this study highlights the potential of metabolic interventions as a strategy for targeted modulation of gene expression programs.

## 1. Introduction

Osteosarcoma is the most common primary malignant bone tumor and is characterized by aggressive local growth, early metastatic dissemination, and poor clinical outcomes in advanced disease (Beird et al., 2022). Although relatively rare, osteosarcoma affects both humans and dogs (Simpson et al., 2017) and represents a major challenge in comparative and translational oncology (Kansara et al., 2014). Current treatment strategies rely primarily on combinations of surgery and chemotherapy; however, survival rates have improved only modestly over recent decades, highlighting the need for novel therapeutic approaches and a deeper understanding of the biological mechanisms underlying tumor progression (Wang et al., 2025).

Cancer development and progression are driven not only by genetic alterations but also by interactions between tumor cells and their microenvironment (De Visser & Joyce, 2023; Mbeunkui & Johann Jr, 2009). Among the many factors shaping cellular behavior, nutrient availability has emerged as a critical regulator of cancer cell physiology (Comito et al. 2020; Lobel et al., 2023). Tumor cells frequently encounter fluctuating levels of nutrients due to abnormal vascularization, rapid proliferation, and competition for resources within the tumor microenvironment. To adapt to these conditions, cancer cells reprogram their metabolism and activate signaling pathways that promote survival, proliferation, and stress resistance. While the metabolic consequences of nutrient limitation have been extensively investigated, the broader effects of nutrient availability on transcriptional regulation remain incompletely understood.

Gene expression is a highly dynamic process that integrates intracellular signaling with environmental stimuli. Changes in nutrient levels can influence transcription through multiple mechanisms, including metabolic signaling pathways, epigenetic modifications, stress-response networks, and growth-factor-dependent regulation (Afman & Muller, 2006; Muller & Kersten, 2003; Sales et al., 2014). Consequently, nutrient availability has the potential to affect not only metabolic genes but also broader biological programs involved in cell fate determination, development, immune regulation, and oncogenesis (Lagoumintzis & Patrinos, 2023). Understanding how nutrient conditions shape gene expression may therefore provide valuable insights into the mechanisms through which the tumor microenvironment influences cancer progression.

Recent advances in high-throughput sequencing technologies have enabled comprehensive characterization of transcriptomic responses to perturbations. In particular, RNA sequencing provides a powerful framework for identifying genes and pathways that respond to specific metabolic conditions (Weglarz-Tomczak et al., 2021a; Weglarz-Tomczak et al., 2021b; Zhang et al., 2025; Zhu et al., 2024). However, most studies have focused on identifying individual differentially expressed genes or enriched pathways, whereas considerably less attention has been devoted to understanding whether nutrient-responsive genes organize into coordinated transcriptional programs. Such programs may reveal higher-order regulatory principles that connect nutrient sensing with complex cellular phenotypes.

In the present study, we investigated the transcriptomic consequences of altering glucose and glutamine availability in an established osteosarcoma cell model (Gill & Gorlick, 2021). Glucose and glutamine were selected because they represent two of the most important metabolic substrates supporting cancer cell growth and biosynthesis (DeBerardinis & Chandel, 2016; Gaglio et al., 2011; Pavlova & Thompson, 2016; Victoria Ruiz-Perez et al., 2013). Using bulk RNA sequencing and differential gene expression analysis, we identified extensive nutrient-dependent transcriptional remodeling across multiple nutrient conditions. Beyond the identification of individual nutrient-responsive genes, our analyses revealed the existence of seven distinct transcriptional programs characterized by highly coordinated expression dynamics. Notably, one of these programs exhibited a strong association with cellular growth and was represented by the proto-oncogene *FOS* (Milde-Langosch, 2005), suggesting a potential link between nutrient availability and growth-associated transcriptional regulation.

Functional annotation further demonstrated that nutrient-responsive genes participate in a wide spectrum of biological processes, including cancer-related pathways, developmental regulation, nervous system function, metabolism, cellular signaling, and immune responses. Together, these findings suggest that nutrient availability acts as a powerful determinant of transcriptomic state and may influence osteosarcoma biology through coordinated regulation of multiple interconnected gene networks.

More broadly, our results support the emerging view that nutrients should not be regarded solely as metabolic substrates, but also as regulators of gene expression. By revealing nutrient-sensitive transcriptional programs in osteosarcoma cells, this study provides a foundation for future investigations into metabolic control of cellular behavior and highlights the potential of nutrient-based interventions as a strategy for targeted modulation of gene expression.

## 2. Results

To investigate how nutrient availability influences gene expression in osteosarcoma cells, we performed a comprehensive transcriptomic analysis of cells cultured under five distinct combinations of glucose and glutamine concentrations, namely, 10mM of glucose and 4mM of glutamine (denoted as Glc10Gln4), 2mM of glucose and 4mM of glutamine (denoted as Glc2Gln4), 25mM of glucose and 4mM of glutamine (denoted as Glc25Gln4), 2mM of glucose and 0.5mM of glutamine (denoted as Glc2Gln0.5), and 2mM of glucose and 2mM of glutamine (denoted as Glc2Gln2); see Table 1. Differential gene expression analysis using DESeq2 (Love et al., 2014; Muzellec et al., 2023) was first used to identify genes whose expression was significantly altered across nutrient conditions. We then examined whether nutrient-responsive genes formed coordinated expression programs and characterized their functional roles using annotations from Gene Ontology (Aleksander et al., 2025; Ashburner et al., 2000). Together, these analyses provide a systematic view of how nutrient availability reshapes the osteosarcoma transcriptome and reveal distinct transcriptional programs associated with cancer-related biological processes.

**Table 1.** The five different media compositions used to culture U2OS cells. * In a form of a stable dipeptide (L-alanine + L-glutamine).

| **Condition Name** | **Glucose (mM)** | **Glutamine (mM)** |
|---|---|---|
| Glc10Gln4 | 10 | 4 |
| Glc2Gln4 | 2 | 4 |
| Glc25Gln4 | 25 | 4* |
| Glc2Gln0.5 | 2 | 0.5 |
| Glc2Gln2 | 2 | 2 |

### 2.1 Nutrient Availability Induces Extensive Transcriptomic Remodeling in Osteosarcoma Cells

#### Global Transcriptomic Response to Nutrient Perturbation

To investigate the impact of nutrient availability on gene expression, we performed bulk RNA sequencing of osteosarcoma cells cultured under five combinations of glucose and glutamine concentrations. Principal component analysis (PCA) was used to assess the global structure of the transcriptomic data and the reproducibility of biological replicates (Figure 1B).

**Figure 1.**
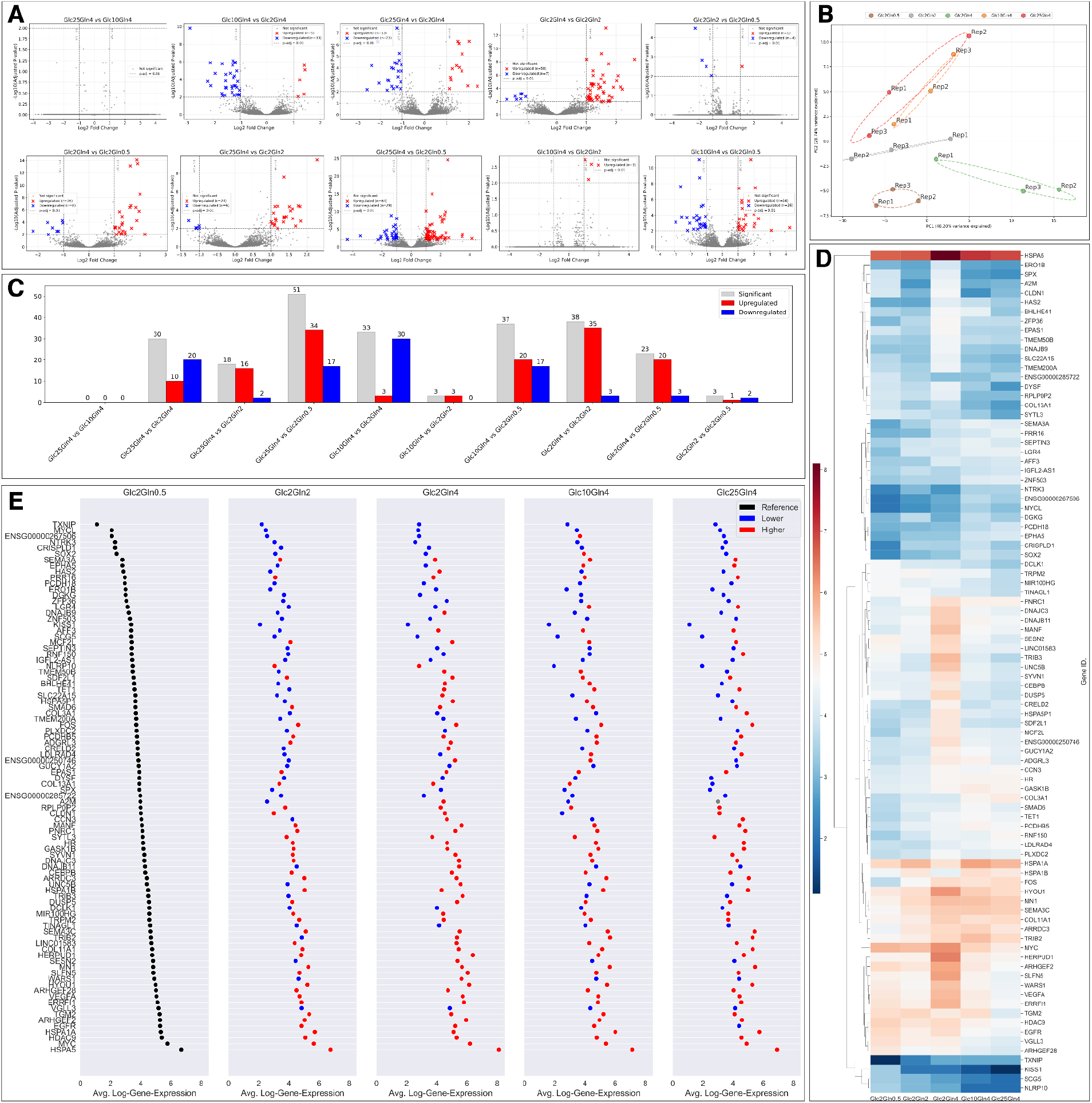
Nutrient availability induces extensive transcriptomic remodeling in osteosarcoma cells. Differential gene expression analysis was performed on osteosarcoma cells cultured under five combinations of glucose and glutamine concentrations. (A) Violin plots summarizing the distribution of differential gene expression statistics for all pairwise comparisons between nutrient conditions. (B) Principal component analysis (PCA) of transcriptomic profiles, showing clustering of biological replicates and separation of nutrient conditions, indicating distinct transcriptional states. (C) Numbers of significantly up- and downregulated genes identified in each comparison. (D) Heatmap of log-transformed gene expression values for all differentially expressed genes across the five nutrient conditions. Hierarchical clustering reveals coordinated groups of genes with similar expression profiles. (E) Relative gene expression profiles using the Glc2Gln0.5 condition as the reference state, highlighting nutrient-dependent up- and downregulation patterns.

The PCA revealed clear separation between nutrient conditions, indicating that alterations in glucose and glutamine availability substantially affected the transcriptional state of the cells. Replicates from the same condition clustered closely together, demonstrating high experimental reproducibility. Moreover, distinct nutrient conditions occupied separate regions of the PCA space, suggesting that nutrient availability drives systematic and condition-specific transcriptomic responses. These results indicate that modulation of extracellular nutrient concentrations is sufficient to induce pronounced global changes in gene expression in osteosarcoma cells.

#### Identification of Differentially Expressed Genes

To identify genes responsive to nutrient perturbation, differential gene expression analysis was performed using DESeq2 (Love et al., 2014) for all pairwise comparisons between the five experimental conditions. The resulting volcano plots are shown in Figure 1A. The extent of transcriptional changes varied substantially across comparisons. While some contrasts yielded only a limited number of significantly differentially expressed genes, others exhibited extensive transcriptional remodeling. In particular, comparisons involving the lowest glutamine concentration (Glc2Gln0.5) frequently produced the largest number of significant genes, suggesting that severe glutamine limitation elicits a strong transcriptional response.

Across all comparisons, both upregulated and downregulated genes were identified, although the relative proportions differed between conditions (Figure 1C). The largest transcriptional differences were observed between conditions associated with markedly different nutrient availability, whereas more moderate changes in nutrient concentrations generally resulted in fewer differentially expressed genes.

Collectively, these findings demonstrate that both glucose and glutamine concentrations influence gene expression in osteosarcoma cells and that the magnitude of the response depends on the specific nutrient environment.

#### Expression Profiles of Nutrient-Responsive Genes

To obtain an overview of the identified nutrient-responsive genes, we visualized their expression patterns across all experimental conditions using hierarchical clustering and heatmap analysis (Figure 1D). The heatmap revealed extensive heterogeneity in gene expression responses, with subsets of genes displaying coordinated increases or decreases across nutrient conditions. Several clusters of genes exhibited similar expression profiles, suggesting the existence of shared regulatory programs. Notably, distinct groups of genes responded preferentially to specific nutrient environments, again indicating that nutrient availability influences multiple transcriptional modules rather than a single global response.

To further examine these trends, gene expression levels were plotted relative to the Glc2Gln0.5 condition, which served as a reference state (Figure 1E). This analysis highlighted genes that were consistently upregulated or downregulated under alternative nutrient conditions and revealed gradual expression changes across the nutrient landscape. The observed patterns suggest that many genes respond quantitatively to nutrient availability rather than displaying simple binary on/off regulation.

Together, these results demonstrate that nutrient perturbation induces widespread and structured transcriptomic remodeling in osteosarcoma cells, motivating a more detailed investigation of coordinated gene expression programs.

### 2.2 Nutrient-Responsive Genes Organize into Distinct Expression Programs

#### Identification of Seven Characteristic Expression Programs

The coordinated expression patterns observed in Figure 1 suggested that nutrient-responsive genes may be governed by shared regulatory mechanisms. To investigate this possibility, we analyzed the expression trajectories of all differentially expressed genes across the five nutrient conditions and grouped genes exhibiting similar responses.

This analysis revealed seven distinct expression programs, each characterized by a unique transcriptional trajectory across the nutrient landscape (Figure 2). Rather than displaying independent gene-specific responses, many genes followed highly similar expression dynamics, indicating the presence of coordinated regulatory modules. The identified programs were designated according to representative genes within each cluster and included the MANF-like (Figure 2.B), FOS-like (Figure 2.C), KISS1-like (Figure 2.D), DGKG-like (Figure 2.E), NTRK3-like (Figure 2.F), EGFR-like (Figure 2.G), and MYCL-like (Figure 2.H) programs.

**Figure 2.**
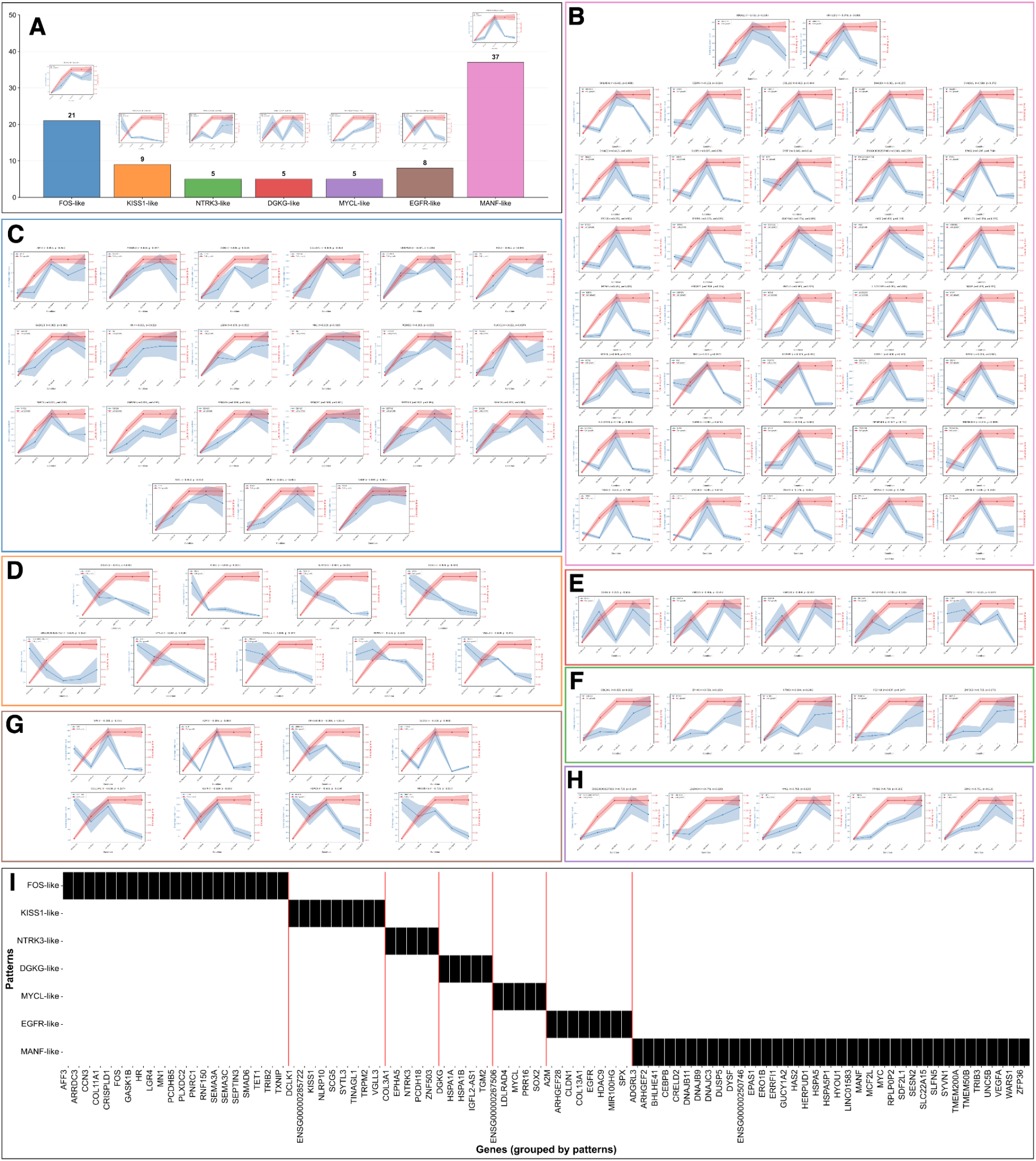
Nutrient-responsive genes organize into seven distinct transcriptional programs. Analysis of expression trajectories across nutrient conditions identified seven characteristic transcriptional programs. (A) Distribution of genes among the identified expression programs. (B–H) Representative expression profiles corresponding to the MANF-like, FOS-like, KISS1-like, DGKG-like, NTRK3-like, EGFR-like, and MYCL-like programs, respectively. Genes within each cluster exhibit highly coordinated responses to changes in glucose and glutamine availability, indicating the presence of shared regulatory mechanisms. The identified programs differ substantially in both their expression dynamics and the number of associated genes. (I) Genes grouped according to their assigned expression programs.

The distribution of genes among the identified programs was highly uneven (Figure 2A). The largest group corresponded to the MANF-like program, comprising 37 genes, followed by the FOS-like program containing 21 genes. The remaining programs contained between 5 and 9 genes each. These findings indicate that nutrient-responsive transcriptional remodeling is concentrated within a limited number of recurrent expression programs rather than being distributed uniformly across individual genes.

#### Characterization of Nutrient-Responsive Expression Programs

Detailed examination of the individual clusters revealed marked differences in how genes responded to changes in nutrient availability (Figure 2B-H). Despite the diversity of responses, genes within each program exhibited remarkable consistency in their expression trajectories, supporting the robustness of the identified classification.

The FOS-like program (Figure 2.C) was characterized by a progressive increase in expression with increasing nutrient availability, followed by a plateau or stabilization under nutrient-rich conditions, achieving highest gene expression in the high glucose and high glutamine levels. However, for a condition which is too high glucose concentration (i.e., above 10), the gene expression level drops. Representative genes within this group included the proto-oncogene *FOS* together with several genes implicated in cellular signaling, proliferation, and stress responses.

In contrast, the MYCL-like (Figure 2.H) and NTRK3-like (Figure 2.F) programs exhibited more selective responses, with expression levels growing with the increasing amount of both glucose and glutamine. The DGKG-like program (Figure 2.E) displayed a distinct biphasic behavior, suggesting sensitivity to specific combinations of glucose and glutamine concentrations. The KISS1-like program (Figure 2.D) expressed a decreasing gene expression level with the growing amount of both glucose and glutamine. The EGFR-like program (Figure 2.G) was very similar to the MYCL-like program but a crucial distinction, namely, much larger gene expression levels for very low amounts of glucose and glutamine (Glc2Gln0.5). The MANF-like program (Figure 2.B) exhibited a pronounced activation profile, peaking at a very particular combination of glucose and glutamine (i.e., Glc2Gln4); moreover, it represented the largest coordinated response identified in the dataset.

Collectively, these observations demonstrate that nutrient availability influences gene expression through multiple distinct regulatory programs rather than through a single global transcriptional mechanism.

#### The FOS-like Program Defines a Growth-Associated Nutrient Response

Among the identified expression programs, the FOS-like cluster emerged as particularly noteworthy. This program contained the proto-oncogene *FOS*, a well-established regulator of cell proliferation, differentiation, stress adaptation, and oncogenic signaling (Milde-Langosch, 2005). Notably, the expression trajectory of genes within this cluster closely mirrored the observed growth characteristics of the osteosarcoma cells across nutrient conditions.

Genes belonging to the FOS-like program exhibited coordinated upregulation under conditions associated with increased cellular growth and metabolic activity, suggesting a close relationship between nutrient availability and proliferation-associated transcriptional responses. The consistency of this pattern across multiple genes further supports the existence of a shared regulatory mechanism linking nutrient sensing to growth-related gene expression.

The identification of a coherent FOS-like transcriptional program is particularly significant because it suggests that nutrient availability may influence osteosarcoma progression not merely through metabolic effects, but also through coordinated activation of growth-associated transcriptional networks. This observation provides a mechanistic framework for understanding how environmental nutrient conditions can shape cellular behavior through large-scale transcriptomic reprogramming.

To facilitate interpretation of the identified programs, all genes were grouped according to their assigned expression patterns (Figure 2I). This organization revealed clear separation between the seven transcriptional programs and highlighted the substantial diversity of nutrient-responsive gene regulation present within the osteosarcoma transcriptome.

### 2.3 Functional Annotation Reveals Broad Biological Roles of Nutrient-Responsive Genes

#### Functional Classification of Nutrient-Responsive Genes

To investigate the biological significance of the identified nutrient-responsive genes, we performed functional annotation using Gene Ontology (GO)-based analysis. The resulting classifications revealed that nutrient-responsive genes participate in a broad range of biological processes associated with cancer, development, nervous system regulation, metabolism, cellular signaling, and immune system function (Figure 3A).

**Figure 3.**
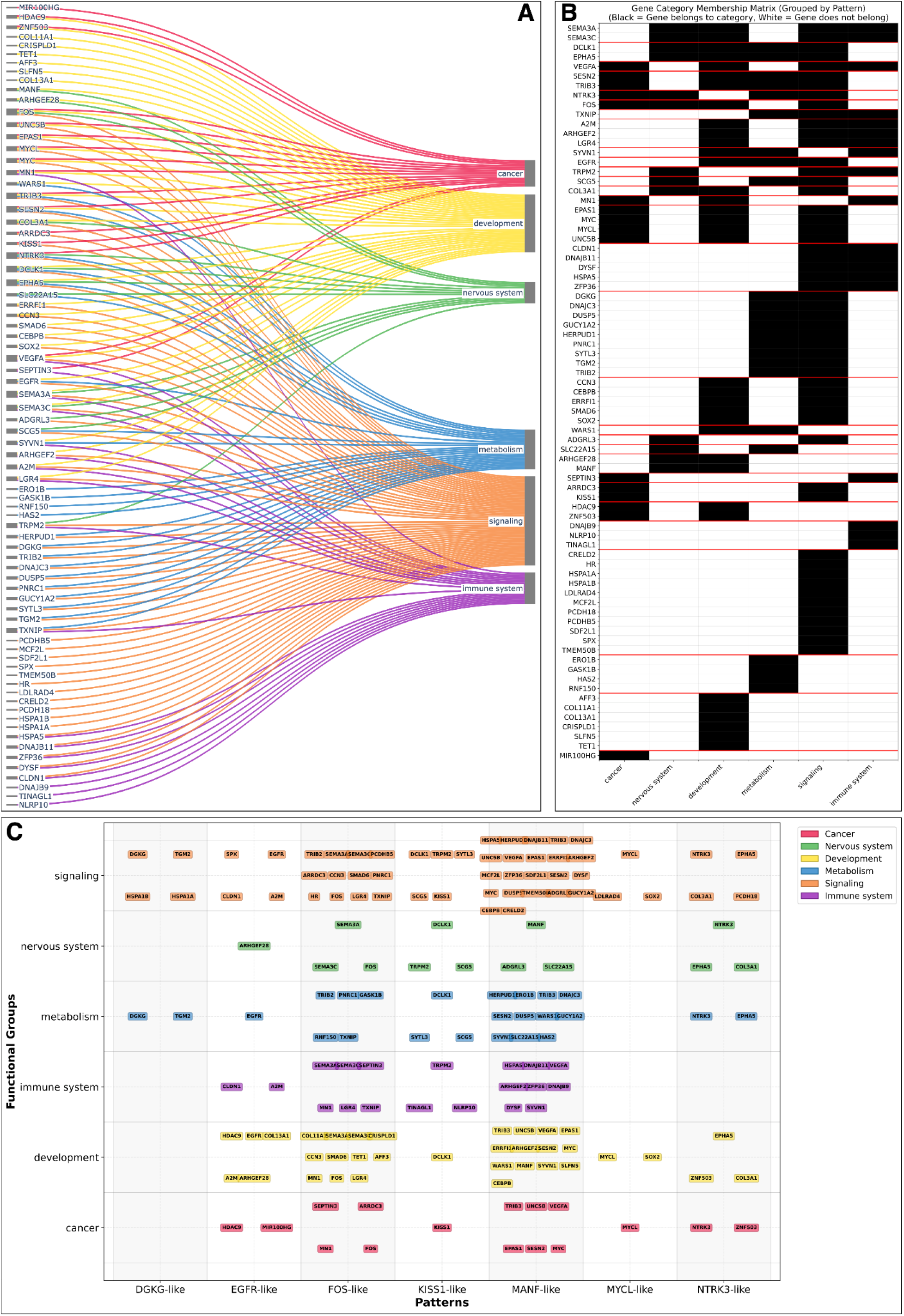
Functional characterization of nutrient-responsive genes reveals broad biological relevance. Gene Ontology-based annotation was used to investigate the biological functions of nutrient-responsive genes. (A) Classification of identified genes into six major functional categories: cancer, nervous system regulation, development, metabolism, signaling, and immune system processes. Many genes participate in multiple biological functions, reflecting the broad impact of nutrient-dependent transcriptional regulation. (B) Network representation illustrating functional overlap between categories and highlighting genes associated with multiple biological processes. (C) Integration of functional annotations with the seven transcriptional programs identified in Figure 2. Genes from all expression programs contribute to multiple biological categories, demonstrating that nutrient-responsive transcriptional remodeling extends across diverse cellular processes. These findings indicate that nutrient availability influences not only metabolic pathways but also gene networks associated with oncogenesis, development, signaling, immunity, and nervous system function.

The identified genes were not confined to a single biological category. Instead, nutrient-responsive transcriptional changes affected multiple functional systems simultaneously, suggesting that nutrient availability acts as a global regulator of cellular state. Genes associated with signaling and metabolic processes constituted a substantial fraction of the identified targets, consistent with the central role of nutrient sensing in coordinating cellular metabolism and growth. However, genes involved in development, immune responses, nervous system regulation, and cancer-related processes were also prominently represented.

Several well-known regulatory genes, including *FOS, MYCL, EGFR, NTRK3, KISS1*, and *MANF*, were among the identified nutrient-responsive genes, further supporting the biological relevance of the observed transcriptional programs.

#### Functional Overlap Between Biological Categories

A detailed examination of the functional annotations revealed extensive overlap between biological categories (Figure 3B). Many genes were assigned to multiple functional groups, indicating that nutrient-responsive genes frequently participate in several biological processes simultaneously. For example, genes associated with cancer-related functions often also appeared within developmental, signaling, or metabolic categories. Similarly, several genes implicated in nervous system regulation were linked to broader cellular signaling pathways. This functional overlap reflects the interconnected nature of biological systems and suggests that nutrient-induced transcriptional changes may propagate through multiple regulatory networks.

The observed pleiotropy further indicates that nutrient availability influences cellular behavior through integrated transcriptional responses rather than through isolated pathway-specific effects. Consequently, modulation of a relatively small number of nutrient-responsive genes may have widespread downstream consequences across diverse biological processes.

#### Integration of Functional Categories with Expression Programs

To better understand the relationship between gene function and transcriptional dynamics, we integrated the functional annotations with the seven expression programs identified in Section 2.2 (Figure 3C).

Genes belonging to all seven transcriptional programs were distributed across multiple functional categories, demonstrating that no individual program was restricted to a single biological process. Nevertheless, distinct trends emerged. The FOS-like program contained genes associated with cancer, development, nervous system regulation, and signaling pathways, consistent with its proposed role in coordinating growth-related responses. Likewise, the KISS1-like, EGFR-like, and NTRK3-like programs contained numerous genes implicated in cellular signaling, developmental regulation, and oncogenic processes. Unlike these programs, the DGKG-like program seems to be more involved in metabolism and signaling, while the MYCL-like program appears to influence cancer, development and signaling.

The MANF-like program, which represented the largest transcriptional cluster, exhibited broad functional diversity and included genes involved in cancer, development, signaling, metabolism, stress responses, and immune-related processes. This observation suggests that the MANF-like program may represent a more general adaptive response to changes in nutrient availability.

Overall, the integration of functional annotations with expression programs revealed that nutrient-responsive transcriptional remodeling extends across multiple biological systems and is organized through distinct yet interconnected regulatory programs. These findings support the notion that nutrient availability influences osteosarcoma biology through coordinated modulation of genes involved in cancer progression, development, metabolism, signaling, immune responses, and nervous system-related processes.

## 3. Discussion

### Nutrient availability acts as a major determinant of transcriptomic state in osteosarcoma cells

The present study demonstrates that relatively small changes in extracellular glucose and glutamine concentrations are sufficient to induce extensive transcriptomic remodeling in an osteosarcoma cell model. Differential gene expression analysis revealed substantial changes across multiple nutrient conditions, while principal component analysis showed clear separation between transcriptomic states associated with distinct nutrient environments. These findings reinforce the long-standing view that nutrients should not be considered solely as metabolic substrates, but also as important regulators of gene expression (Cousins, 1999). Although metabolic adaptation to nutrient availability has been extensively studied, our results highlight the breadth of transcriptional responses that accompany such adaptations and suggest that nutrient sensing may exert widespread control over cellular behavior through gene regulatory mechanisms.

### A central finding of this study is the identification of seven distinct nutrient-responsive transcriptional programs

Rather than responding independently, nutrient-sensitive genes organized into a limited number of highly coordinated expression patterns characterized by reproducible transcriptional trajectories across nutrient conditions. This observation suggests the existence of shared regulatory mechanisms that couple nutrient availability to specific gene expression programs. Importantly, the identified programs displayed markedly different expression dynamics, indicating that nutrient-dependent regulation is not governed by a single global response but instead involves multiple regulatory modules operating in parallel. Such organization may enable cells to selectively activate distinct biological processes depending on the metabolic context.

### Among the identified programs, the FOS-like signature emerged as particularly noteworthy in osteosarcoma

The proto-oncogene *FOS* is a well-established regulator of proliferation, differentiation, stress responses, and oncogenic signaling (Milde-Langosch, 2005) . In our study, genes belonging to the FOS-like program exhibited expression patterns that closely mirrored cellular growth characteristics across nutrient conditions, suggesting a potential link between nutrient availability and growth-associated transcriptional regulation. This finding is consistent with previous observations that immediate-early response genes integrate environmental and metabolic signals to coordinate adaptive cellular responses. The coordinated behavior of multiple genes within this program further supports the hypothesis that nutrient availability may influence osteosarcoma progression through activation of specific transcriptional networks rather than solely through direct metabolic effects.

### The functional diversity of nutrient-responsive genes underscores the broad biological impact of nutrient-dependent transcriptional regulation

Functional annotation revealed associations with cancer-related pathways, developmental processes, nervous system regulation, metabolism, cellular signaling, and immune responses. Notably, many genes participated in multiple functional categories, reflecting the interconnected nature of biological regulatory networks. This observation suggests that nutrient-induced changes in gene expression may propagate across multiple cellular systems simultaneously. Consequently, modulation of nutrient availability has the potential to influence a wide range of biological processes extending well beyond metabolic adaptation.

### The relationship between nutrient availability and cancer biology deserves particular attention

Many of the identified genes, including *FOS, MYCL, EGFR, NTRK3*, and *KISS1*, have previously been implicated in tumor growth, cellular differentiation, metastasis, or therapeutic response. The observation that expression of these genes can be substantially altered through changes in nutrient conditions raises the intriguing possibility that aspects of tumor behavior may be influenced through metabolic interventions. While nutrient modulation alone is unlikely to replace established cancer therapies, our findings suggest that manipulation of the metabolic environment could represent a complementary strategy for influencing cancer-associated transcriptional programs.

### Beyond oncology, the identified transcriptional responses may have broader biological significance

The enrichment of genes associated with development, nervous system regulation, and immune function suggests that nutrient-responsive gene regulation is not restricted to cancer-specific pathways. Instead, nutrient availability may represent a general mechanism through which cells coordinate diverse biological processes in response to environmental conditions. This perspective aligns with growing evidence linking metabolism to developmental decisions, immune regulation, tissue homeostasis, and cellular plasticity.

### Several limitations of the present study should be acknowledged

First, the analyses were performed using a single osteosarcoma cell model, and the extent to which the identified transcriptional programs generalize across additional osteosarcoma models or other cancer types remains to be determined. Second, the study focused on transcriptomic responses and did not directly investigate the upstream regulatory mechanisms responsible for the observed expression patterns. Third, although the identified transcriptional programs exhibit strong biological coherence, functional validation will be required to establish their causal roles in cellular growth and adaptation. Future studies incorporating additional cell lines, perturbation experiments, and mechanistic analyses will be important for further elucidating the regulatory networks underlying nutrient-responsive gene expression.

### Taken together, our findings support a model in which nutrient availability functions as a powerful regulator of transcriptomic organization

By identifying seven distinct nutrient-responsive transcriptional programs and linking them to diverse biological processes, this study provides new insights into how metabolic conditions shape gene expression landscapes in osteosarcoma cells. These observations contribute to a growing appreciation of the intimate relationship between metabolism and gene regulation and establish a foundation for future investigations into nutrient-dependent control of cellular behavior.

## 4. Conclusions

In this study, we demonstrated that nutrient availability profoundly influences gene expression in osteosarcoma cells. By systematically varying glucose and glutamine concentrations and analyzing transcriptomic responses using RNA sequencing, we identified extensive nutrient-dependent transcriptional remodeling involving genes associated with cancer, development, nervous system regulation, metabolism, signaling, and immune function.

Beyond the identification of individual differentially expressed genes, our analyses revealed seven distinct nutrient-responsive transcriptional programs characterized by coordinated expression dynamics across nutrient conditions. Among these, a FOS-like program exhibited a striking association with cellular growth, suggesting a potential mechanistic link between nutrient availability and growth-associated transcriptional regulation. These findings indicate that nutrient-responsive genes are organized into higher-order regulatory modules rather than acting as isolated transcriptional units.

Collectively, our results support a model in which nutrients function not only as metabolic substrates but also as regulators of transcriptomic organization. This perspective opens new opportunities for understanding how environmental and metabolic factors shape cellular behavior in cancer and other biological contexts. More broadly, the ability to modulate gene expression through controlled changes in nutrient availability raises the possibility of developing novel metabolic strategies for influencing cellular function. Ultimately, these findings suggest that nutrients may represent a previously underappreciated class of regulators capable of reshaping transcriptomic states and influencing cellular behavior across a broad range of biological systems.

## 5. Materials and methods

### 5.1 Culturing U2OS Cancer Cells

#### Culturing U2OS Cancer Cells

U2OS (human osteosarcoma) cells were cultured in Dulbecco’s Modified Eagle Medium (DMEM; GibcoBRL, Grand Island, NY, USA), supplemented with 10% Fetal Bovine Serum (FBS) and 1% antibiotics (penicillin and streptomycin) at various initial concentrations of glucose and glutamine/L-alanine-glutamine, see Table 1.

Four of the compositions were made using Advanced DMEM that does not contain glucose, glutamine, phenol red or sodium pyruvate. Glc25Gln4 is commercially available (DMEM glutamaxTM). The cells were first cultured in DMEM glutamaxTM for two weeks until a sufficient number of cells was acquired. Then cells were split over five experiments and cultured for three weeks to adopt.

#### Growth Rate/Cell Proliferation Assay

The total number of cells in the consequent supernatant was determined by hemocytometer counting. Mean growth rate was determined by counting cells in four non-overlapping sets of sixteen corner squares selected at random.

#### Isolation and Measurement of the Metabolites from Medium

Samples from each medium were collected at 0 and 48 h, repeated three times for a total of three independent experiments for every medium composition. Metabolites were isolated using the protocol described in (Sapcariu et al., 2014) with minor modifications. The samples were filtered using 0.22μm syringe filters (BGB Analytik Vertrieb GmbH, Rheinfelden, Germany), freeze dried and stored until measurement at -80°C. Standard solutions for glutamine, glucose and lactate were prepared in the same way as well as control samples. High-performance liquid chromatography (HPLC) was performed using the HPLC-DAD RID LC-20AT Prominence (Shimadzu, Columbia, SC, USA) machine with a UV Diode Array Detector SPD-M30A NexeraX2 or/and a Refractive Index Detector RID 20A and an analytical ion-exclusion Rezex ROA-Organic Acid H+(8%) column (250 × 4.6 mm) with guard column (Phenomenex, Torrance, CA, USA) (5 mM H2SO4 in MilliQ water (18.2 MΩ), isocratic, 0.15 mL/min. flow rate). Injection volume was 15 μL (Autosampler: SIL-20AC, Prominence, Shimadzu), column oven temperature was 55°C (Column oven: CTO-20A, Prominence, Shimadzu) and the pressure was 29 bar.

#### Isolation and Obtaining of RNA Data

A total of 5,000,000 cells from each medium condition were taken for RNA analysis, also repeated three times for a total of three independent experiments for every cell condition. The RNA extraction was done using the TRI Reagent Protocol from Sigma Aldrich^1^, after which all samples were treated with DNase using the TURBO DNA-free Kit from Invitrogen. Nanodrop was used for determining RNA concentrations when needed. With the Agilent 2100 Bioanalyzer System the quality of the RNA was checked, as well as the RNA Integrity Number (RIN), using RNA Pico Chips. rRNA depletion and library preparation for sequencing was completed with the NEBNext protocol (‘Protocol for use with NEBNext rRNA Depletion Kit (Human/Mouse/Rat) (NEB #E6310) and NEBNext Ultra II Directional RNA Library Prep Kit for Illumina (NEB #E7760, #E7765)’) from New England BioLabs. Using a 2200 TapeStation System with Agilent D1000 ScreenTapes (Agilent Technologies, Santa Clara, CA, USA ), and the size distribution of the libraries with indexed adapters was assessed. Quantification of the libraries was performed on a QuantStudio 3 Real-Time PCR System (Thermo Fisher Scientific, Waltham, MA, USA), which was done using the NEBNext Library Quant Kit for Illumina (New England BioLabs, Ipswich, MA, USA) according to the manufacturer’s directions. On a NextSeq 550 Sequencing System (Illumina), the libraries were clustered and sequenced (75 bp) utilizing a NextSeq 500/550 High Output Kit v2.5 (75 Cycles) (Illumina).

#### RNA-Seq Data Processing

RNA-seq data were available in .fastq file format. There were three measurements per condition. The RNA-seq data were processed according to the following pipeline:

- Sequences were mapped against the human reference genome using Burrows-Wheeler Aligner^2^. The outputs of this step were files in .sam format.
- The .sam files were transformed to sorted bam files using Samtools^3^.
- Gene occurrences were calculated using featuresCounts from the Subread package^4^.
- In all cases, the Homo sapiens (human) genome assembly GRCh38 (hg38) from Genome Reference Consortium Build 38^5^ was used as the reference genome

### 5.2 Differential Gene Expression Analysis using DESeq2

#### Differential Gene Expression Analysis

Bulk RNA-seq data comprising 67,149 genes across 5 experimental conditions (n=3 biological replicates per condition) were analyzed for differential gene expression using PyDESeq2 v0.4.12 (Muzellec et al., 2023), the Python implementation of the DESeq2 algorithm (Love et al., 2014). Raw count matrices were quality-controlled and filtered to remove low-abundance transcripts prior to analysis.

#### Statistical Analysis

Differential expression analysis was performed using the DESeq2 framework with default parameters. The analysis pipeline included: (1) estimation of size factors for library normalization using the median-of-ratios method, (2) estimation of gene-wise dispersions using empirical Bayes shrinkage, and (3) fitting of generalized linear models with negative binomial distribution to model count data. The experimental design matrix specified condition as the primary factor of interest.

#### Pairwise Comparisons

All possible pairwise comparisons between the 5 experimental conditions were performed using Wald tests, resulting in 10 total comparisons. For each comparison, differential expression statistics including log_2_ fold changes, Wald statistics, and p-values were calculated. Multiple testing correction was applied using the Benjamini-Hochberg false discovery rate (FDR) method.

#### Significance Thresholds

Genes were considered significantly differentially expressed if they met the following criteria: adjusted p-value (padj) < 0.05 and absolute log_2_ fold change > 1.0, corresponding to a minimum 2-fold change in expression. These thresholds were selected to balance statistical significance with biological relevance.

#### Data Processing and Normalization

Size factor-normalized counts were used for downstream visualization and analysis. Principal component analysis (PCA) was performed on variance-stabilized transformed data to assess sample clustering and identify potential batch effects. All statistical analyses were performed in Python using the PyDESeq2 package, with results exported for visualization and functional enrichment analysis.

#### Functional Classification of Differentially Expressed Genes

Significantly differentially expressed genes were systematically classified into functional categories based on Gene Ontology (GO)^6^ (Aleksander et al., 2025; Ashburner et al., 2000) annotations and established biological keyword searches. A comprehensive classification framework was developed targeting six major functional categories: cancer-related processes, nervous system function, immune system processes, developmental biology, metabolism, and cellular signaling pathways.

The classification algorithm integrated multiple annotation sources for each gene, including: (1) gene symbol and name annotations, (2) GO molecular function, biological process, and cellular component terms, (3) pathway annotations, and (4) functional descriptions. For each functional category, curated keyword lists and relevant GO terms were established based on domain expertise and literature review.

Functional Categories and Criteria:

- **Cancer-related**: Keywords encompassing oncogenes, tumor suppressors, cell cycle regulation, apoptosis, DNA repair, and proliferation processes
- **Nervous System**: Terms related to neuronal development, synaptic function, neurotransmitter systems, and brain-specific processes
- **Immune System**: Immune response components including cytokines, immune cell types, inflammatory processes, and antigen presentation
- **Development**: Embryonic development, cell differentiation, stem cell biology, and organogenesis processes
- **Metabolism**: Enzymatic activities, metabolic pathways, and energy production processes
- **Signaling**: Signal transduction pathways, receptors, kinases, and second messenger systems

Gene classification was performed by text-based matching against gene annotations, with genes assigned to functional categories based on keyword presence in combined annotation text (gene descriptions, GO terms, and pathway annotations). Genes could be assigned to multiple categories if annotations supported diverse functional roles.

## 6. Supplementary material

More detailed results and more details about genes can be found in Supplementary Material.

## Footnotes

1 https://www.sigmaaldrich.com/technical-documents/protocols/biology/tri-reagent.html

2 BWA; version 0.7.17, https://sourceforge.net/projects/bio-bwa/

3 https://anaconda.org/bioconda/samtools

4 https://anaconda.org/bioconda/subread

5 https://www.ncbi.nlm.nih.gov/assembly/GCF_000001405.26/

6 https://geneontology.org/

## Notes

### Competing Interest Statement

The authors have declared no competing interest.

