## Supplementary Material for "Nutrient Availability as a Driver of Transcriptome Reprogramming in Osteosarcoma"

### Nutrient Availability as a Driver of Transcriptome Reprogramming in Osteosarcoma
#### *Supplementary Material*

**Jakub M. Tomczak**[1], **Zuzanna Senkowska**[1], **Ewelina Węglarz-Tomczak**[1]

[1] GenSiLab Inc.

### Differential Gene Expression Analysis (DESeq2): The complete list of up- and downregulated genes

In the following, we present detailed results of differential gene expression analysis using DESeq2 between two conditions. In the tables, the following columns are shown:

- **Gene ID** represented by ENSEMBL ID;
- **baseMean** representing a mean of gene expression levels from three measurements;
- **log2FoldChange** represents the effect size estimate of how much a gene's expression has changed between two conditions;
- **lfcSE** corresponding to Log Fold Change Standard Error;
- **stat** indicating the value of the Wald test;
- **pvalue** being the p-value;
- **padj** being the adjusted p-value.

##### Glc2Gln2 vs Glc2Gln0.5

| Gene ID. | baseMean | log2FoldChange | lfcSE | stat | pvalue | padj |
|---|---|---|---|---|---|---|
| ENSG00000163347 | 36.759 | -1.798 | 0.345 | -5.209 | 0.000 | 0.001 |
| ENSG00000175899 | 32.861 | -2.280 | 0.378 | -6.036 | 0.000 | 0.000 |
| ENSG00000170345 | 131.310 | 1.132 | 0.229 | 4.943 | 0.000 | 0.003 |

##### Glc2Gln4 vs Glc2Gln0.5

| Gene ID. | baseMean | log2FoldChange | lfcSE | stat | pvalue | padj |
|---|---|---|---|---|---|---|
| ENSG00000142910 | 65.422 | -1.033 | 0.200 | -5.171 | 0.000 | 0.000 |

| Gene ID. | baseMean | log2FoldChange | lfcSE | stat | pvalue | padj |
|---|---|---|---|---|---|---|
| ENSG00000215895 | 61.363 | 1.648 | 0.255 | 6.471 | 0.000 | 0.000 |
| ENSG00000145050 | 108.699 | 1.872 | 0.211 | 8.890 | 0.000 | 0.000 |
| ENSG00000090520 | 107.576 | 1.286 | 0.256 | 5.023 | 0.000 | 0.000 |
| ENSG00000150471 | 74.607 | 1.318 | 0.295 | 4.461 | 0.000 | 0.003 |
| ENSG00000250746 | 75.615 | 1.611 | 0.254 | 6.345 | 0.000 | 0.000 |
| ENSG00000146278 | 110.369 | 1.195 | 0.198 | 6.035 | 0.000 | 0.000 |
| ENSG00000075213 | 44.195 | 1.281 | 0.314 | 4.084 | 0.000 | 0.009 |
| ENSG00000128590 | 32.854 | 1.427 | 0.328 | 4.348 | 0.000 | 0.004 |
| ENSG00000170961 | 30.581 | 1.673 | 0.356 | 4.704 | 0.000 | 0.001 |
| ENSG00000044574 | 1292.581 | 1.628 | 0.191 | 8.513 | 0.000 | 0.000 |
| ENSG00000162298 | 79.281 | 1.033 | 0.215 | 4.795 | 0.000 | 0.001 |
| ENSG00000152402 | 71.433 | 1.042 | 0.220 | 4.733 | 0.000 | 0.001 |
| ENSG00000149428 | 217.504 | 1.293 | 0.226 | 5.714 | 0.000 | 0.000 |
| ENSG00000133083 | 51.751 | -1.199 | 0.270 | -4.440 | 0.000 | 0.003 |
| ENSG00000102580 | 95.111 | 1.308 | 0.253 | 5.172 | 0.000 | 0.000 |
| ENSG00000126217 | 67.335 | 2.000 | 0.280 | 7.151 | 0.000 | 0.000 |
| ENSG00000170345 | 131.310 | 1.902 | 0.220 | 8.653 | 0.000 | 0.000 |
| ENSG00000051108 | 186.012 | 1.870 | 0.348 | 5.375 | 0.000 | 0.000 |
| ENSG00000128016 | 43.246 | 1.960 | 0.322 | 6.086 | 0.000 | 0.000 |
| ENSG00000128228 | 56.700 | 1.836 | 0.290 | 6.324 | 0.000 | 0.000 |
| ENSG00000285722 | 30.751 | -1.629 | 0.311 | -5.234 | 0.000 | 0.000 |
| ENSG00000142188 | 40.158 | 1.076 | 0.266 | 4.052 | 0.000 | 0.010 |

#### Glc2Gln4 vs Glc2Gln2

| Gene ID. | baseMean | log2FoldChange | lfcSE | stat | pvalue | padj |
|---|---|---|---|---|---|---|
| ENSG00000116285 | 151.824 | 1.215 | 0.188 | 6.458 | 0.000 | 0.000 |
| ENSG00000130766 | 103.423 | 1.123 | 0.238 | 4.725 | 0.000 | 0.001 |
| ENSG00000215895 | 61.363 | 1.738 | 0.254 | 6.837 | 0.000 | 0.000 |
| ENSG00000163393 | 33.257 | 1.427 | 0.325 | 4.389 | 0.000 | 0.002 |
| ENSG00000116584 | 174.349 | 1.076 | 0.234 | 4.602 | 0.000 | 0.001 |
| ENSG00000086619 | 20.086 | 1.492 | 0.376 | 3.967 | 0.000 | 0.009 |
| ENSG00000116016 | 46.253 | 1.490 | 0.252 | 5.916 | 0.000 | 0.000 |
| ENSG00000145050 | 108.699 | 1.507 | 0.202 | 7.446 | 0.000 | 0.000 |
| ENSG00000058866 | 26.850 | -1.363 | 0.326 | -4.184 | 0.000 | 0.005 |
| ENSG00000090520 | 107.576 | 1.079 | 0.252 | 4.272 | 0.000 | 0.004 |
| ENSG00000163347 | 36.759 | 2.066 | 0.338 | 6.111 | 0.000 | 0.000 |
| ENSG00000250746 | 75.615 | 1.552 | 0.251 | 6.192 | 0.000 | 0.000 |
| ENSG00000204389 | 268.506 | -1.047 | 0.234 | -4.479 | 0.000 | 0.002 |
| ENSG00000204388 | 122.965 | -1.181 | 0.248 | -4.761 | 0.000 | 0.001 |
| ENSG00000112715 | 143.018 | 1.246 | 0.227 | 5.498 | 0.000 | 0.000 |
| ENSG00000164484 | 37.444 | 1.196 | 0.294 | 4.060 | 0.000 | 0.007 |
| ENSG00000128590 | 32.854 | 1.583 | 0.330 | 4.803 | 0.000 | 0.001 |
| ENSG00000170961 | 30.581 | 1.898 | 0.360 | 5.269 | 0.000 | 0.000 |
| ENSG00000044574 | 1292.581 | 1.651 | 0.191 | 8.638 | 0.000 | 0.000 |
| ENSG00000162298 | 79.281 | 1.200 | 0.217 | 5.542 | 0.000 | 0.000 |
| ENSG00000152402 | 71.433 | 1.190 | 0.221 | 5.387 | 0.000 | 0.000 |

| Gene ID. | baseMean | log2FoldChange | lfcSE | stat | pvalue | padj |
|---|---|---|---|---|---|---|
| ENSG00000149428 | 217.504 | 1.126 | 0.225 | 5.014 | 0.000 | 0.000 |
| ENSG00000107731 | 90.162 | 2.147 | 0.404 | 5.314 | 0.000 | 0.000 |
| ENSG00000138166 | 85.077 | 1.403 | 0.270 | 5.189 | 0.000 | 0.000 |
| ENSG00000175899 | 32.861 | 2.621 | 0.370 | 7.089 | 0.000 | 0.000 |
| ENSG00000134548 | 29.356 | 1.849 | 0.422 | 4.384 | 0.000 | 0.002 |
| ENSG00000123095 | 45.331 | 1.599 | 0.282 | 5.665 | 0.000 | 0.000 |
| ENSG00000102580 | 95.111 | 1.394 | 0.253 | 5.511 | 0.000 | 0.000 |
| ENSG00000126217 | 67.335 | 1.106 | 0.257 | 4.309 | 0.000 | 0.003 |
| ENSG00000140105 | 127.143 | 1.217 | 0.198 | 6.143 | 0.000 | 0.000 |
| ENSG00000259518 | 98.225 | 1.153 | 0.277 | 4.159 | 0.000 | 0.005 |
| ENSG00000051108 | 186.012 | 2.018 | 0.348 | 5.792 | 0.000 | 0.000 |
| ENSG00000166750 | 151.838 | 1.758 | 0.388 | 4.535 | 0.000 | 0.001 |
| ENSG00000101255 | 94.126 | 2.200 | 0.431 | 5.103 | 0.000 | 0.000 |
| ENSG00000128016 | 43.246 | 1.316 | 0.299 | 4.395 | 0.000 | 0.002 |
| ENSG00000128228 | 56.700 | 1.474 | 0.280 | 5.272 | 0.000 | 0.000 |
| ENSG00000184164 | 54.766 | 1.356 | 0.295 | 4.597 | 0.000 | 0.001 |
| ENSG00000142188 | 40.158 | 1.450 | 0.273 | 5.304 | 0.000 | 0.000 |

#### Glc10Gln4 vs Glc2Gln0.5

| Gene ID. | baseMean | log2FoldChange | lfcSE | stat | pvalue | padj |
|---|---|---|---|---|---|---|
| ENSG00000142910 | 65.422 | -1.054 | 0.204 | -5.179 | 0.000 | 0.000 |
| ENSG00000116990 | 16.370 | 1.995 | 0.456 | 4.372 | 0.000 | 0.003 |

| Gene ID. | baseMean | log2FoldChange | lfcSE | stat | pvalue | padj |
|---|---|---|---|---|---|---|
| ENSG00000265972 | 11.033 | 3.085 | 0.762 | 4.048 | 0.000 | 0.009 |
| ENSG00000170498 | 8.921 | -2.912 | 0.614 | -4.747 | 0.000 | 0.001 |
| ENSG00000071575 | 175.030 | 1.232 | 0.200 | 6.148 | 0.000 | 0.000 |
| ENSG00000168542 | 79.415 | 1.319 | 0.326 | 4.050 | 0.000 | 0.009 |
| ENSG00000206538 | 108.718 | -1.444 | 0.179 | -8.088 | 0.000 | 0.000 |
| ENSG00000181449 | 27.220 | 2.040 | 0.439 | 4.644 | 0.000 | 0.001 |
| ENSG00000163347 | 36.759 | -2.598 | 0.378 | -6.880 | 0.000 | 0.000 |
| ENSG00000150471 | 74.607 | 1.239 | 0.297 | 4.171 | 0.000 | 0.006 |
| ENSG00000145242 | 32.247 | 1.466 | 0.323 | 4.545 | 0.000 | 0.002 |
| ENSG00000214944 | 90.328 | -1.420 | 0.195 | -7.276 | 0.000 | 0.000 |
| ENSG00000113369 | 147.723 | 1.236 | 0.206 | 6.004 | 0.000 | 0.000 |
| ENSG00000184838 | 41.727 | 1.359 | 0.314 | 4.331 | 0.000 | 0.003 |
| ENSG00000113209 | 75.438 | 1.263 | 0.222 | 5.680 | 0.000 | 0.000 |
| ENSG00000164674 | 35.649 | -1.369 | 0.319 | -4.291 | 0.000 | 0.004 |
| ENSG00000048052 | 152.465 | -1.211 | 0.206 | -5.874 | 0.000 | 0.000 |
| ENSG00000146648 | 133.012 | -1.320 | 0.228 | -5.796 | 0.000 | 0.000 |
| ENSG00000075223 | 188.050 | 1.051 | 0.156 | 6.756 | 0.000 | 0.000 |
| ENSG00000075213 | 44.195 | 2.078 | 0.305 | 6.802 | 0.000 | 0.000 |
| ENSG00000121005 | 25.415 | 1.985 | 0.371 | 5.352 | 0.000 | 0.000 |
| ENSG00000182261 | 15.122 | -2.538 | 0.494 | -5.140 | 0.000 | 0.000 |
| ENSG00000205213 | 51.423 | 1.470 | 0.278 | 5.296 | 0.000 | 0.000 |
| ENSG00000243742 | 39.313 | -1.591 | 0.303 | -5.247 | 0.000 | 0.000 |

| Gene ID. | baseMean | log2FoldChange | lfcSE | stat | pvalue | padj |
|---|---|---|---|---|---|---|
| ENSG00000255248 | 64.100 | -1.052 | 0.231 | -4.547 | 0.000 | 0.002 |
| ENSG00000138336 | 64.859 | 1.336 | 0.269 | 4.958 | 0.000 | 0.000 |
| ENSG00000197467 | 27.517 | -1.598 | 0.328 | -4.866 | 0.000 | 0.000 |
| ENSG00000175899 | 32.861 | -1.771 | 0.348 | -5.086 | 0.000 | 0.000 |
| ENSG00000134548 | 29.356 | -2.217 | 0.434 | -5.107 | 0.000 | 0.000 |
| ENSG00000133083 | 51.751 | -1.405 | 0.277 | -5.074 | 0.000 | 0.000 |
| ENSG00000170345 | 131.310 | 1.776 | 0.222 | 8.010 | 0.000 | 0.000 |
| ENSG00000166922 | 15.355 | -2.184 | 0.540 | -4.042 | 0.000 | 0.009 |
| ENSG00000137834 | 60.777 | 1.084 | 0.217 | 4.988 | 0.000 | 0.000 |
| ENSG00000267506 | 19.353 | 2.271 | 0.453 | 5.016 | 0.000 | 0.000 |
| ENSG00000169184 | 207.319 | 1.013 | 0.236 | 4.300 | 0.000 | 0.004 |
| ENSG00000100167 | 48.230 | 1.125 | 0.255 | 4.410 | 0.000 | 0.003 |
| ENSG00000285722 | 30.751 | -1.425 | 0.310 | -4.599 | 0.000 | 0.001 |

#### Glc10Gln4 vs Glc2Gln2

| Gene ID. | baseMean | log2FoldChange | lfcSE | stat | pvalue | padj |
|---|---|---|---|---|---|---|
| ENSG00000071575 | 175.030 | 1.081 | 0.198 | 5.467 | 0.000 | 0.002 |
| ENSG00000075213 | 44.195 | 1.320 | 0.269 | 4.901 | 0.000 | 0.008 |
| ENSG00000123095 | 45.331 | 1.506 | 0.285 | 5.284 | 0.000 | 0.003 |

#### Glc10Gln4 vs Glc2Gln4

| Gene ID. | baseMean | log2FoldChange | lfcSE | stat | pvalue | padj |
|---|---|---|---|---|---|---|
| ENSG00000116285 | 151.824 | -1.166 | 0.186 | -6.282 | 0.000 | 0.000 |
| ENSG00000130766 | 103.423 | -1.120 | 0.235 | -4.759 | 0.000 | 0.001 |
| ENSG00000215895 | 61.363 | -1.167 | 0.239 | -4.875 | 0.000 | 0.001 |
| ENSG00000163393 | 33.257 | -1.519 | 0.324 | -4.696 | 0.000 | 0.001 |
| ENSG00000116584 | 174.349 | -1.250 | 0.234 | -5.353 | 0.000 | 0.000 |
| ENSG00000086619 | 20.086 | -1.571 | 0.374 | -4.200 | 0.000 | 0.006 |
| ENSG00000116016 | 46.253 | -1.371 | 0.245 | -5.603 | 0.000 | 0.000 |
| ENSG00000135636 | 36.784 | -1.311 | 0.271 | -4.831 | 0.000 | 0.001 |
| ENSG00000145050 | 108.699 | -1.267 | 0.197 | -6.435 | 0.000 | 0.000 |
| ENSG00000058866 | 26.850 | 1.385 | 0.322 | 4.297 | 0.000 | 0.005 |
| ENSG00000163347 | 36.759 | -2.865 | 0.371 | -7.723 | 0.000 | 0.000 |
| ENSG00000250746 | 75.615 | -1.020 | 0.239 | -4.264 | 0.000 | 0.005 |
| ENSG00000204389 | 268.506 | 1.397 | 0.232 | 6.024 | 0.000 | 0.000 |
| ENSG00000204388 | 122.965 | 1.420 | 0.245 | 5.793 | 0.000 | 0.000 |
| ENSG00000112715 | 143.018 | -1.016 | 0.223 | -4.565 | 0.000 | 0.002 |
| ENSG00000164484 | 37.444 | -1.296 | 0.293 | -4.421 | 0.000 | 0.003 |
| ENSG00000128590 | 32.854 | -1.540 | 0.324 | -4.754 | 0.000 | 0.001 |
| ENSG00000044574 | 1292.581 | -1.192 | 0.190 | -6.264 | 0.000 | 0.000 |
| ENSG00000243742 | 39.313 | -1.543 | 0.298 | -5.175 | 0.000 | 0.000 |
| ENSG00000162298 | 79.281 | -1.088 | 0.212 | -5.128 | 0.000 | 0.000 |
| ENSG00000107731 | 90.162 | -1.652 | 0.396 | -4.169 | 0.000 | 0.007 |
| ENSG00000138166 | 85.077 | -1.665 | 0.271 | -6.131 | 0.000 | 0.000 |

| Gene ID. | baseMean | log2FoldChange | lfcSE | stat | pvalue | padj |
|---|---|---|---|---|---|---|
| ENSG00000175899 | 32.861 | -2.113 | 0.340 | -6.221 | 0.000 | 0.000 |
| ENSG00000134548 | 29.356 | -2.207 | 0.430 | -5.135 | 0.000 | 0.000 |
| ENSG00000102580 | 95.111 | -1.214 | 0.248 | -4.890 | 0.000 | 0.001 |
| ENSG00000140105 | 127.143 | -1.118 | 0.195 | -5.736 | 0.000 | 0.000 |
| ENSG00000259518 | 98.225 | -1.296 | 0.277 | -4.683 | 0.000 | 0.001 |
| ENSG00000051108 | 186.012 | -1.937 | 0.347 | -5.589 | 0.000 | 0.000 |
| ENSG00000166750 | 151.838 | -1.675 | 0.386 | -4.341 | 0.000 | 0.004 |
| ENSG00000101255 | 94.126 | -2.097 | 0.428 | -4.901 | 0.000 | 0.001 |
| ENSG00000172216 | 69.295 | -1.203 | 0.221 | -5.444 | 0.000 | 0.000 |
| ENSG00000128228 | 56.700 | -1.534 | 0.278 | -5.527 | 0.000 | 0.000 |
| ENSG00000184164 | 54.766 | -1.214 | 0.289 | -4.205 | 0.000 | 0.006 |

#### Glc25Gln4 vs Glc2Gln0.5

| Gene ID. | baseMean | log2FoldChange | lfcSE | stat | pvalue | padj |
|---|---|---|---|---|---|---|
| ENSG00000142910 | 65.422 | -1.039 | 0.217 | -4.788 | 0.000 | 0.000 |
| ENSG00000116990 | 16.370 | 2.053 | 0.467 | 4.393 | 0.000 | 0.002 |
| ENSG00000060718 | 165.839 | 1.143 | 0.214 | 5.330 | 0.000 | 0.000 |
| ENSG00000265972 | 11.033 | 3.470 | 0.766 | 4.528 | 0.000 | 0.001 |
| ENSG00000170498 | 8.921 | -3.459 | 0.749 | -4.620 | 0.000 | 0.001 |
| ENSG00000071575 | 175.030 | 1.262 | 0.204 | 6.196 | 0.000 | 0.000 |
| ENSG00000135636 | 36.784 | -1.826 | 0.341 | -5.359 | 0.000 | 0.000 |
| ENSG00000144218 | 41.973 | 1.214 | 0.282 | 4.308 | 0.000 | 0.003 |

| Gene ID. | baseMean | log2FoldChange | lfcSE | stat | pvalue | padj |
|---|---|---|---|---|---|---|
| ENSG00000168542 | 79.415 | 2.221 | 0.324 | 6.853 | 0.000 | 0.000 |
| ENSG00000206538 | 108.718 | -1.355 | 0.188 | -7.208 | 0.000 | 0.000 |
| ENSG00000181449 | 27.220 | 2.108 | 0.447 | 4.715 | 0.000 | 0.001 |
| ENSG00000145242 | 32.247 | 2.104 | 0.322 | 6.535 | 0.000 | 0.000 |
| ENSG00000189184 | 27.734 | 1.595 | 0.340 | 4.693 | 0.000 | 0.001 |
| ENSG00000170153 | 66.686 | 2.101 | 0.310 | 6.768 | 0.000 | 0.000 |
| ENSG00000164125 | 96.399 | 1.094 | 0.246 | 4.441 | 0.000 | 0.002 |
| ENSG00000214944 | 90.328 | -1.214 | 0.203 | -5.991 | 0.000 | 0.000 |
| ENSG00000113369 | 147.723 | 1.036 | 0.212 | 4.893 | 0.000 | 0.000 |
| ENSG00000184838 | 41.727 | 2.345 | 0.309 | 7.586 | 0.000 | 0.000 |
| ENSG00000113209 | 75.438 | 1.433 | 0.227 | 6.308 | 0.000 | 0.000 |
| ENSG00000204388 | 122.965 | 1.000 | 0.247 | 4.048 | 0.000 | 0.006 |
| ENSG00000146278 | 110.369 | 1.258 | 0.204 | 6.163 | 0.000 | 0.000 |
| ENSG00000164674 | 35.649 | -1.741 | 0.353 | -4.936 | 0.000 | 0.000 |
| ENSG00000048052 | 152.465 | -1.041 | 0.211 | -4.926 | 0.000 | 0.000 |
| ENSG00000146648 | 133.012 | -1.063 | 0.232 | -4.576 | 0.000 | 0.001 |
| ENSG00000075223 | 188.050 | 1.423 | 0.157 | 9.052 | 0.000 | 0.000 |
| ENSG00000075213 | 44.195 | 2.266 | 0.310 | 7.301 | 0.000 | 0.000 |
| ENSG00000168453 | 87.251 | 1.098 | 0.237 | 4.638 | 0.000 | 0.001 |
| ENSG00000121005 | 25.415 | 1.674 | 0.389 | 4.307 | 0.000 | 0.003 |
| ENSG00000136999 | 81.603 | 1.141 | 0.257 | 4.448 | 0.000 | 0.002 |
| ENSG00000136997 | 263.908 | -1.032 | 0.180 | -5.721 | 0.000 | 0.000 |

| Gene ID. | baseMean | log2FoldChange | lfcSE | stat | pvalue | padj |
|---|---|---|---|---|---|---|
| ENSG00000182261 | 15.122 | -2.090 | 0.500 | -4.177 | 0.000 | 0.004 |
| ENSG00000205213 | 51.423 | 1.984 | 0.279 | 7.117 | 0.000 | 0.000 |
| ENSG00000255248 | 64.100 | -1.011 | 0.243 | -4.152 | 0.000 | 0.004 |
| ENSG00000120594 | 61.134 | 1.190 | 0.295 | 4.040 | 0.000 | 0.006 |
| ENSG00000138336 | 64.859 | 1.476 | 0.275 | 5.374 | 0.000 | 0.000 |
| ENSG00000197467 | 27.517 | -1.666 | 0.356 | -4.676 | 0.000 | 0.001 |
| ENSG00000165655 | 42.225 | 1.602 | 0.308 | 5.207 | 0.000 | 0.000 |
| ENSG00000134548 | 29.356 | -2.059 | 0.454 | -4.538 | 0.000 | 0.001 |
| ENSG00000133083 | 51.751 | -1.657 | 0.299 | -5.544 | 0.000 | 0.000 |
| ENSG00000126217 | 67.335 | 1.448 | 0.292 | 4.953 | 0.000 | 0.000 |
| ENSG00000170345 | 131.310 | 2.496 | 0.221 | 11.287 | 0.000 | 0.000 |
| ENSG00000166922 | 15.355 | -2.262 | 0.582 | -3.885 | 0.000 | 0.010 |
| ENSG00000137834 | 60.777 | 1.046 | 0.226 | 4.624 | 0.000 | 0.001 |
| ENSG00000140538 | 19.033 | 1.987 | 0.465 | 4.276 | 0.000 | 0.003 |
| ENSG00000267506 | 19.353 | 2.373 | 0.461 | 5.145 | 0.000 | 0.000 |
| ENSG00000168675 | 64.339 | 1.446 | 0.323 | 4.476 | 0.000 | 0.001 |
| ENSG00000198959 | 147.390 | -1.302 | 0.280 | -4.659 | 0.000 | 0.001 |
| ENSG00000268621 | 39.962 | 1.098 | 0.264 | 4.160 | 0.000 | 0.004 |
| ENSG00000169184 | 207.319 | 1.268 | 0.237 | 5.338 | 0.000 | 0.000 |
| ENSG00000100167 | 48.230 | 1.113 | 0.265 | 4.207 | 0.000 | 0.003 |
| ENSG00000142185 | 76.394 | -1.011 | 0.237 | -4.263 | 0.000 | 0.003 |

#### Glc25Gln4 vs Glc2Gln2

| Gene ID. | baseMean | log2FoldChange | lfcSE | stat | pvalue | padj |
|---|---|---|---|---|---|---|
| ENSG00000060718 | 165.839 | 1.194 | 0.214 | 5.574 | 0.000 | 0.000 |
| ENSG00000071575 | 175.030 | 1.111 | 0.201 | 5.526 | 0.000 | 0.000 |
| ENSG00000144218 | 41.973 | 1.368 | 0.283 | 4.836 | 0.000 | 0.001 |
| ENSG00000168542 | 79.415 | 1.768 | 0.315 | 5.620 | 0.000 | 0.000 |
| ENSG00000145242 | 32.247 | 1.541 | 0.293 | 5.261 | 0.000 | 0.000 |
| ENSG00000189184 | 27.734 | 1.608 | 0.336 | 4.789 | 0.000 | 0.001 |
| ENSG00000170153 | 66.686 | 1.598 | 0.297 | 5.375 | 0.000 | 0.000 |
| ENSG00000164125 | 96.399 | 1.136 | 0.246 | 4.626 | 0.000 | 0.001 |
| ENSG00000184838 | 41.727 | 2.289 | 0.301 | 7.596 | 0.000 | 0.000 |
| ENSG00000075213 | 44.195 | 1.509 | 0.275 | 5.488 | 0.000 | 0.000 |
| ENSG00000168453 | 87.251 | 1.119 | 0.236 | 4.752 | 0.000 | 0.001 |
| ENSG00000136999 | 81.603 | 1.090 | 0.254 | 4.287 | 0.000 | 0.005 |
| ENSG00000120594 | 61.134 | 1.185 | 0.293 | 4.048 | 0.000 | 0.010 |
| ENSG00000165655 | 42.225 | 1.351 | 0.298 | 4.540 | 0.000 | 0.002 |
| ENSG00000170345 | 131.310 | 1.363 | 0.197 | 6.917 | 0.000 | 0.000 |
| ENSG00000168675 | 64.339 | 1.810 | 0.328 | 5.516 | 0.000 | 0.000 |
| ENSG00000198959 | 147.390 | -1.291 | 0.279 | -4.624 | 0.000 | 0.001 |
| ENSG00000142185 | 76.394 | -1.008 | 0.236 | -4.266 | 0.000 | 0.005 |

#### Glc25Gln4 vs Glc2Gln4

| Gene ID. | baseMean | log2FoldChange | lfcSE | stat | pvalue | padj |
|---|---|---|---|---|---|---|

| | | | | | | |
|---|---|---|---|---|---|---|
| ENSG00000116285 | 151.824 | -1.140 | 0.193 | -5.916 | 0.000 | 0.000 |
| ENSG00000130766 | 103.423 | -1.197 | 0.246 | -4.874 | 0.000 | 0.001 |
| ENSG00000116584 | 174.349 | -1.335 | 0.240 | -5.556 | 0.000 | 0.000 |
| ENSG00000135636 | 36.784 | -2.105 | 0.332 | -6.333 | 0.000 | 0.000 |
| ENSG00000168542 | 79.415 | 2.014 | 0.313 | 6.436 | 0.000 | 0.000 |
| ENSG00000145050 | 108.699 | -1.036 | 0.202 | -5.129 | 0.000 | 0.000 |
| ENSG00000163347 | 36.759 | -1.397 | 0.329 | -4.241 | 0.000 | 0.006 |
| ENSG00000145242 | 32.247 | 1.696 | 0.289 | 5.879 | 0.000 | 0.000 |
| ENSG00000189184 | 27.734 | 1.685 | 0.328 | 5.135 | 0.000 | 0.000 |
| ENSG00000184838 | 41.727 | 1.442 | 0.268 | 5.390 | 0.000 | 0.000 |
| ENSG00000204389 | 268.506 | 1.497 | 0.234 | 6.406 | 0.000 | 0.000 |
| ENSG00000204388 | 122.965 | 1.554 | 0.248 | 6.252 | 0.000 | 0.000 |
| ENSG00000112715 | 143.018 | -1.183 | 0.231 | -5.118 | 0.000 | 0.000 |
| ENSG00000136997 | 263.908 | -1.245 | 0.178 | -6.992 | 0.000 | 0.000 |
| ENSG00000044574 | 1292.581 | -1.042 | 0.191 | -5.457 | 0.000 | 0.000 |
| ENSG00000205213 | 51.423 | 1.178 | 0.243 | 4.838 | 0.000 | 0.001 |
| ENSG00000162298 | 79.281 | -1.246 | 0.226 | -5.513 | 0.000 | 0.000 |
| ENSG00000107731 | 90.162 | -1.673 | 0.404 | -4.147 | 0.000 | 0.008 |
| ENSG00000165655 | 42.225 | 1.568 | 0.295 | 5.313 | 0.000 | 0.000 |
| ENSG00000138166 | 85.077 | -1.477 | 0.279 | -5.293 | 0.000 | 0.000 |
| ENSG00000175899 | 32.861 | -1.599 | 0.342 | -4.680 | 0.000 | 0.001 |
| ENSG00000134548 | 29.356 | -2.049 | 0.450 | -4.557 | 0.000 | 0.002 |
| ENSG00000123095 | 45.331 | -1.239 | 0.285 | -4.352 | 0.000 | 0.004 |

| | | | | | | |
|---|---|---|---|---|---|---|
| ENSG00000102580 | 95.111 | -1.075 | 0.255 | -4.222 | 0.000 | 0.006 |
| ENSG00000140105 | 127.143 | -1.150 | 0.203 | -5.652 | 0.000 | 0.000 |
| ENSG00000140538 | 19.033 | 1.954 | 0.441 | 4.428 | 0.000 | 0.003 |
| ENSG00000051108 | 186.012 | -1.651 | 0.349 | -4.731 | 0.000 | 0.001 |
| ENSG00000101255 | 94.126 | -2.377 | 0.440 | -5.403 | 0.000 | 0.000 |
| ENSG00000172216 | 69.295 | -1.065 | 0.231 | -4.615 | 0.000 | 0.002 |
| ENSG00000268621 | 39.962 | 1.204 | 0.255 | 4.717 | 0.000 | 0.001 |

##### Glc25Gln4 vs Glc10Gln4

None found

### Main representatives of the seven distinct expression programs

#### MANF

Mesencephalic astrocyte-derived neurotrophic factor (*MANF*) is an endoplasmic reticulum (ER) stress-inducible protein characterized by two main structural domains: a saposin-like lipid-binding domain at the N-terminus and a C-terminal SAP domain, joined by a short linker region. In humans, MANF is encoded by a gene that is located on the short arm of chromosome 3. After cleavage of the signal peptide, MANF is secreted as a mature protein consisting of 158 amino acid residues with a molecular weight of approximately 18 kDa. Initially identified for its neuroprotective functions, *MANF* plays a critical role in maintaining cellular homeostasis under stress conditions. Beyond the nervous system, *MANF* has shown cytoprotective effects in several non-neuronal disease models, such as retinal degeneration, myocardial infarction, liver injury, nephrotic syndrome, and diabetes mellitus (Liu et al., 2022). It is particularly important for pancreatic β-cell health, where its absence results in prolonged unfolded protein response activation and impaired cell viability (Matlik et al., 2015). Additionally, *MANF* has anti-inflammatory properties. It inhibits NF-κB signaling by preventing the p65 transcription factor from binding to the promoter regions of its target genes, thus modulating immune and inflammatory responses (Chen et al., 2015).

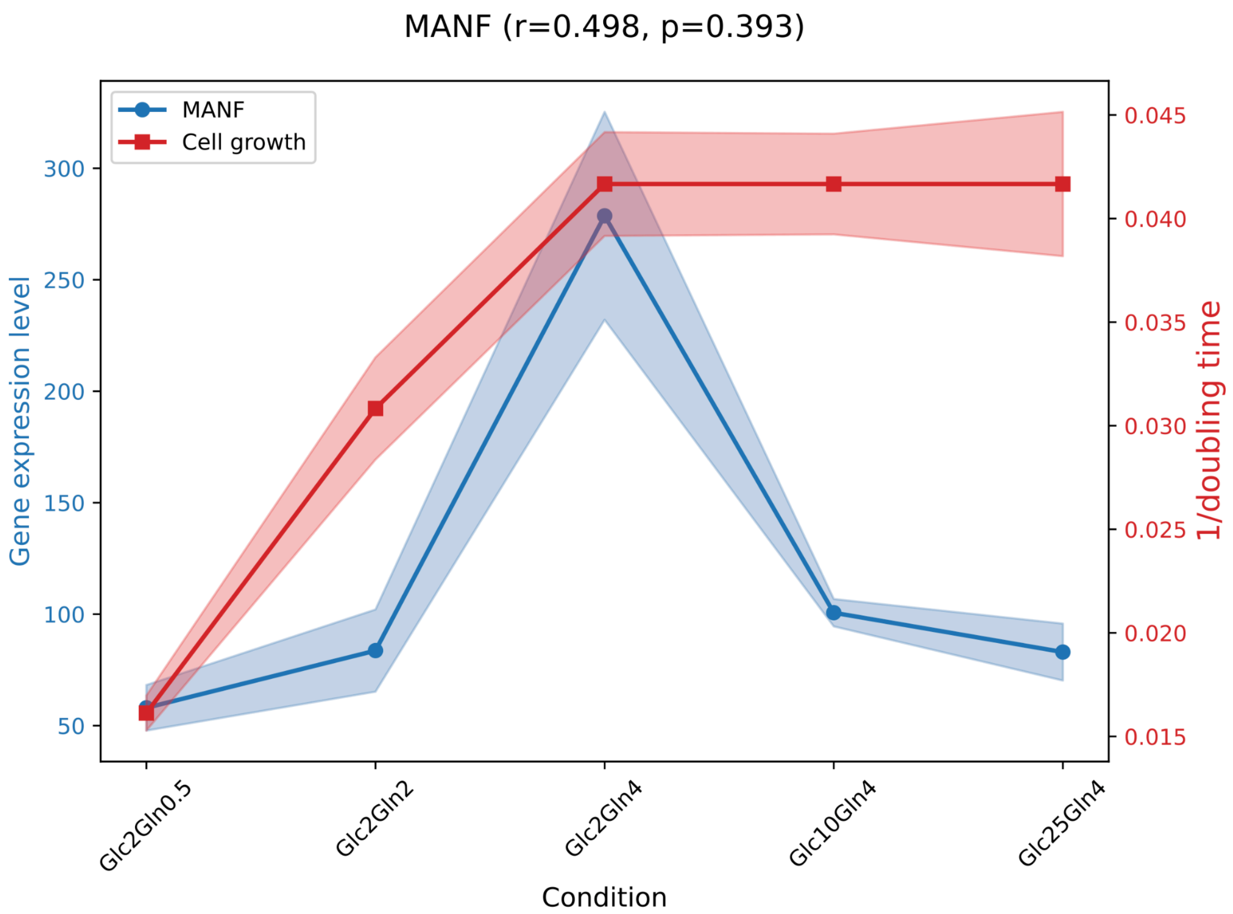


**Figure S1. The gene expression for *MANF* under various conditions.**

### FOS

FOS is a nuclear proto-oncogene expressed in a wide range of tissues, encoded in humans on chromosome 14. It is the cellular homolog of the oncogenes carried by the FBJ and FBR murine osteogenic sarcoma viruses. The *FOS* gene is part of a multigene family that includes FOSB, FRA-1, and FRA-2, all of which contain a leucine-rich "leucine zipper" domain involved in protein-protein interactions, and a basic region required for DNA binding. Members of the *FOS* family, including c-Fos, can form heterodimers with Jun family proteins, leading to the formation of the AP-1 transcription factor complex (Johnson et al., 1992). AP-1 regulates transcription of various genes involved in cell proliferation and cancer development (Xu et al., 2022). Several genes such as those encoding collagenase, interleukin-2, and adipocyte P2 contain AP-1 binding sites as cis-acting transcriptional regulatory sequences (Johnson et al., 1992). The *FOS* gene encodes a 380-amino-acid protein with a molecular weight of 56-62 kDa. The transactivation domain is located at the C-terminus (Dalhausser et al., 2022). Activation of c-Fos is modulated by several transcriptional regulators including calcium ($Ca^{2+}$), cyclic AMP (cAMP), and mitogen-activated protein kinases (MAPKs) (Lara Aparicio et al., 2022). It plays an essential role in cellular processes such as growth, osteoclastogenesis, hematopoiesis, neuronal activation, excitability, and survival (Dalhausser et al., 2022)

As an immediate-early response gene, *FOS* is rapidly upregulated in response to numerous extracellular stimuli including nutritional signals, oxidative stress, and growth factors like epidermal growth factor (EGF) (Manimala et al., 2013). In human hepatocellular carcinoma cells, EGF stimulates FOS expression, promoting hepatocyte proliferation and hepatocellular carcinoma development. In colon cancer, the phosphorylated (active) form of the c-Fos protein, via the cytokine GDF15, caused an increase in tumor metastasis and invasiveness (Ding, et al., 2020)

Interestingly, c-Fos also exhibits self-regulatory effects. The c-Fos protein can negatively regulate its own expression, potentially enabling a rapid shutdown of IEG (immediate early gene) activity after stimulus. This repression requires ongoing protein synthesis (Lara Aparicio et al., 2022).

In cancer research, c-Fos is known to influence drug resistance. Along with other oncogenes such as H-ras and K-ras, c-Fos regulates expression levels of genes encoding efflux pumps, DNA repair enzymes, and detoxifying enzymes. roles confirmed through ribozyme experiments (Clynes et al., 1998). A study conducted on the adriamycin (ADR)-selected human breast cancer cell line with the multidrug resistance phenotype (MCF-7/ADR) showed that silencing of the FOS gene sensitized cancer cells to chemotherapy. FOS downregulation resulted in decreased expression and activity of P-glycoprotein (P-gp; transmembrane protein which facilitates unilateral transports of the intracellular drugs out of the cells contributing to cell resistance), enhanced apoptosis, and altered expression of apoptosis-associated proteins such as Bax, Bcl-2, p53, and PUMA) (Shi et al., 2013) Another study examined the effect of c-Fos on resistance to 5-fluorouracil (5-FU) in colorectal cancer. Elevated levels of c-Fos were shown to promote resistance to 5-FU by enhancing cancer stem cell characteristics through the direct regulation of the pluripotency-related transcription factor NANOG. In contrast, reducing c-Fos expression increased the sensitivity of resistant cells to 5-FU. Overall, these findings indicate that targeting c-Fos activity could be a potential therapeutic approach for overcoming drug resistance in colon cancer (Gui, et al., 2024).

Experimental studies have shown that cardiac hormones can significantly downregulate *FOS*, reducing its levels by 59%-61% in hepatocellular cancer cells, and by 74%-82% in small-cell lung cancer and renal adenocarcinoma cells (Xu et al., 2022).

In neuroscience, *FOS* expression is often used as a marker for neuronal activation. Its expression is rapidly induced by various stimuli and is applied to investigate how external signals affect nerve cell phenotype. However, the specific role of c-Fos proteins in the nervous system remains to be fully elucidated. Expression patterns depend on the type of stimulus, brain region, and downstream signaling mechanisms (Lara Aparicio et al., 2022).

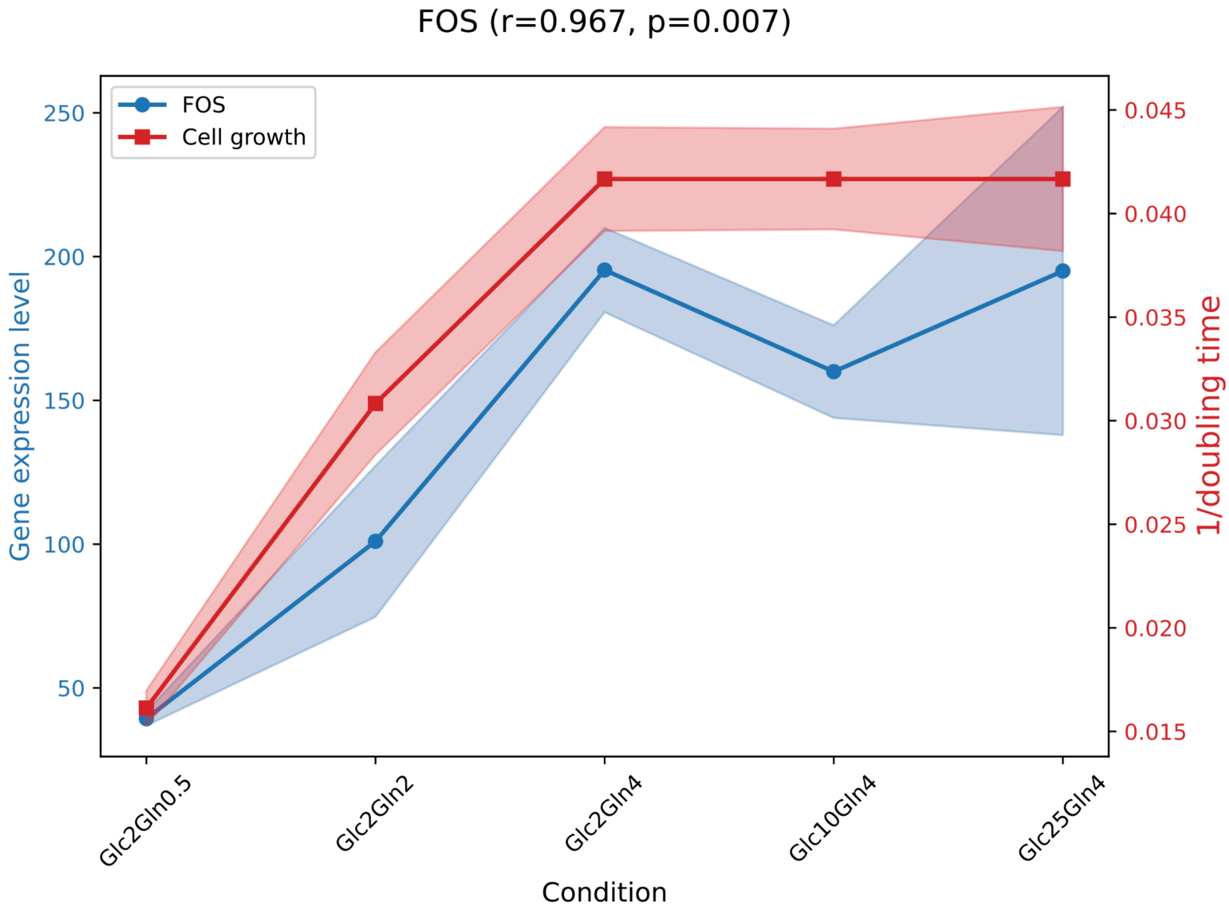


**Figure S2. The gene expression for *FOS* under various conditions.**

### KISS1

Kisspeptin (KSS), the protein product of the KISS1 gene, was initially identified as a metastasis suppressor in human malignant melanoma cells and was later shown to inhibit breast cancer metastasis as well. The human KISS1 gene, located on chromosome 1, encodes a pre-proprotein that is cleaved into several bioactive kisspeptin peptides, all of which activate the KISS1R receptor. Despite being functionally conserved, the amino acid sequences of kisspeptin show relatively low similarity across species, with only 50-52% sequence homology between humans and rodents, and even lower similarity with nonmammalian species. In humans, kisspeptin is most abundantly expressed in the placenta, followed by the pancreas, liver, and small intestine. During pregnancy, maternal levels of kisspeptin rise dramatically, by several thousand-fold, with the placenta serving as the main source. It plays key roles during gestation, including supporting embryo implantation, placental development, and regulation of maternal glucose metabolism (Aljubouri et al., 2025; Fratangelo, et al., 2018).

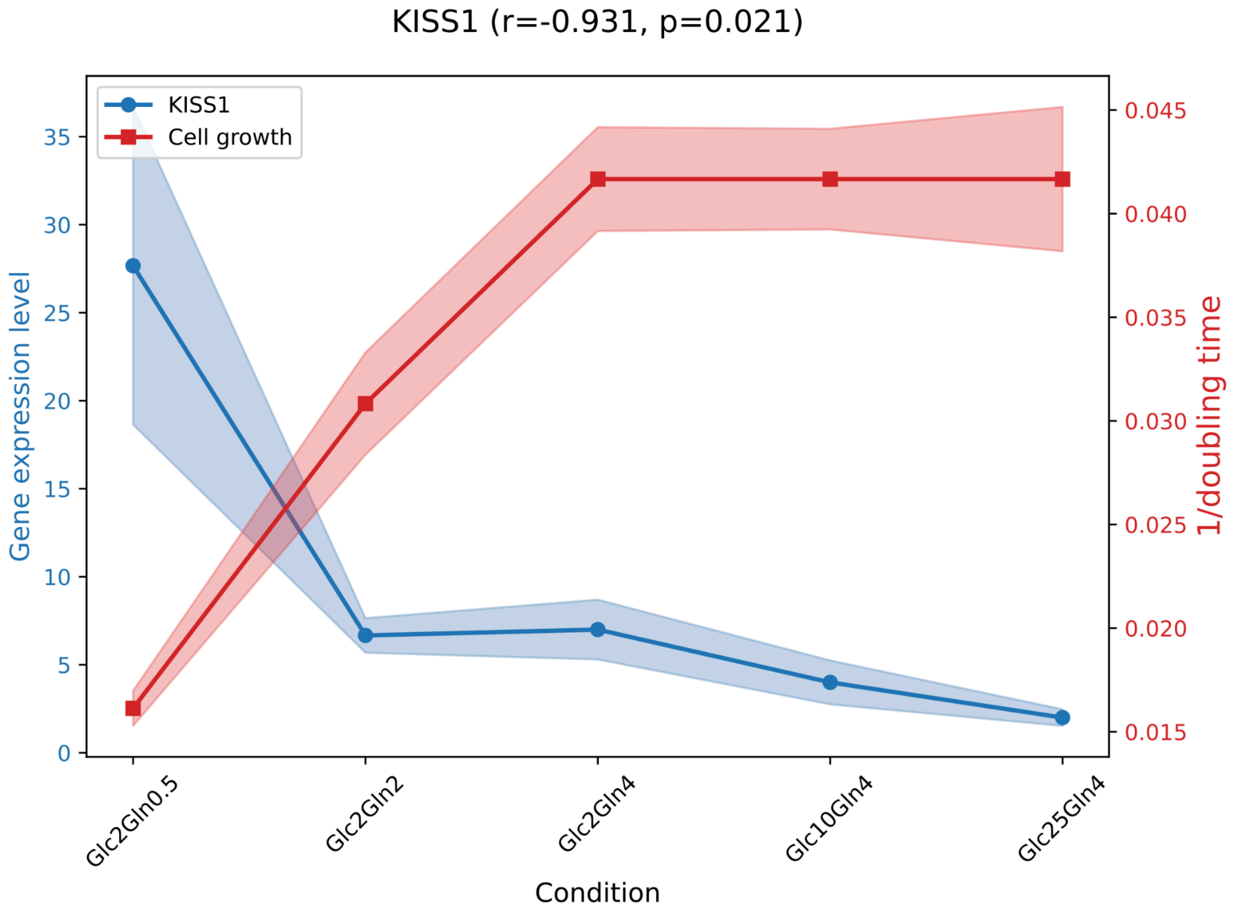


**Figure S3. The gene expression for *KISS1* under various conditions.**

### DGKG

Diacylglycerol kinase (DGK) gene is localized on the long arm of chromosome 3. Its protein product catalyzes the phosphorylation of diacylglycerol (DG) to generate phosphatidic acid (PA), leading to an activation of protein kinase C (PKC). PA also serves as an activator for several other kinases, including phosphatidylinositol 4-phosphate 5-kinase, and the mammalian target of rapamycin (mTOR). As such, DGK is considered a crucial component in lipid-mediated signal transduction pathways. Studies involving DGK knockout mice have underscored its vital functions in the immune system, cardiac physiology, insulin resistance associated with diabetes, and brain function (Barber & Raben, 2020). This evolutionarily conserved enzyme family exists across all multicellular organisms (Luo et al., 2004). DGKG is classified as a type I DGK, characterized by the presence of EF-hand motifs and a recovering homology (RVH) domain which makes them calcium-dependent. It contains two cysteine-rich regions (C1a and C1b) in its N-terminal regulatory domain and a catalytic domain at the C-terminus with molecular weight of 90kDa. DGKG may influence macrophage differentiation by modulating the expression of key genes, such as those encoding adhesion molecules and lysosomal enzymes. Additionally, DGKG might function as a negative regulator of macrophage differentiation in leukemia cells, specifically, through catalytic activity affecting the cytoskeleton (Yamada et al., 2003).

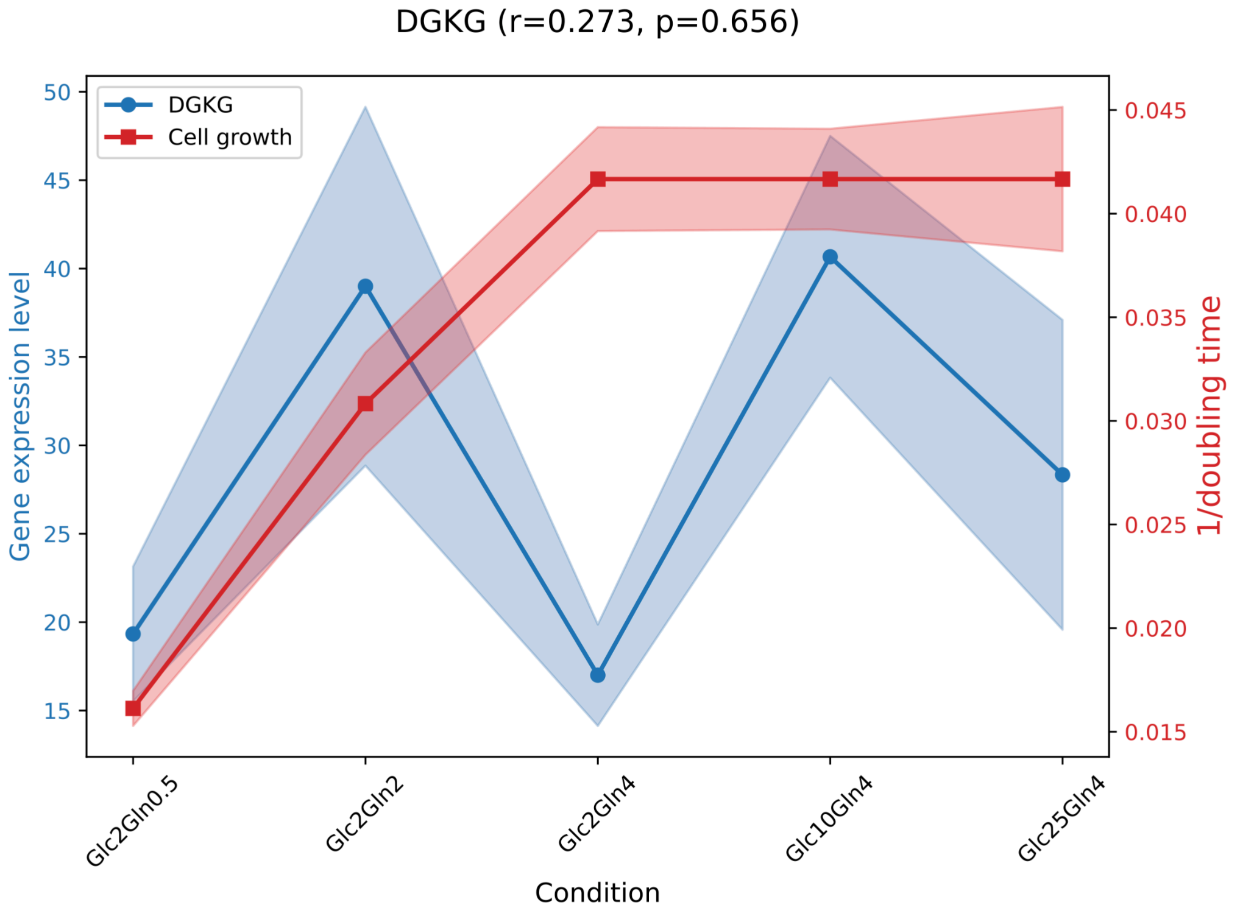


**Figure S4. The gene expression for *DGKG* under various conditions.**

#### NTRK3

The neurotrophic receptor tyrosine kinase 3 (NTRK3) gene is located on chromosome 15q25.3 and encodes TrkC, a receptor tyrosine kinase with a molecular weight of approximately 94 kDa, that primarily binds neurotrophin-3 (NT-3). Upon ligand binding, TrkC activates signaling pathways such as PI3K-AKT, RAS-MAPK, and PLCγ, regulating neuronal survival, differentiation, proliferation, and synaptic plasticity. NTRK3 plays a crucial role in the development and maintenance of the nervous system.

Genetic alterations involving NTRK3, particularly gene fusions, have been identified in several cancers (Manea et al., 2022), including secretory breast carcinoma, secretory salivary gland carcinoma, and various pediatric and adult solid tumors. The ETV6-NTRK3 fusion is the best-characterized alteration and results in constitutive kinase activation that drives tumorigenesis (Kinnunen et al., 2023). Tumors harboring NTRK3 fusions can be effectively treated with selective TRK inhibitors such as larotrectinib and entrectinib, making NTRK3 an important target in precision oncology (Hagopian & Nagasaka, 2024).

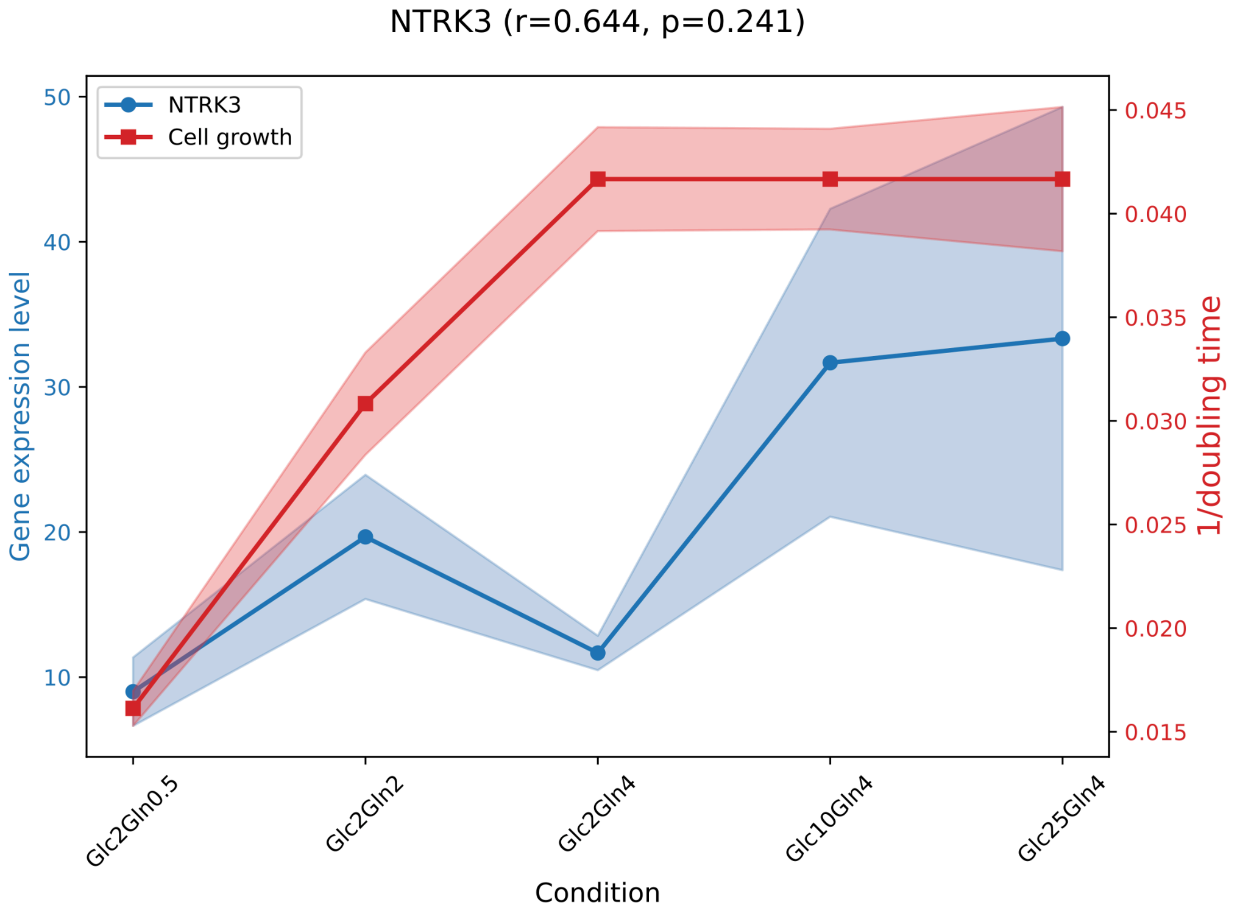


**Figure S5. The gene expression for *NTRK3* under various conditions.**

### EGFR

The epidermal growth factor receptor (EGFR) gene is located on chromosome 7p11.2 and encodes a receptor tyrosine kinase belonging to the ERBB family, with a molecular weight of approximately 170 kDa. Upon binding growth factors such as EGF, EGFR activates signaling pathways including RAS-MAPK, PI3K-AKT, and JAK-STAT, thereby regulating cell proliferation, survival, differentiation, and migration. EGFR is essential for normal epithelial tissue development and homeostasis.

Aberrant EGFR signaling caused by gene amplification, overexpression, or activating mutations contributes to the development of numerous cancers (Sigismund et al., 2018), including non-small cell lung cancer (NSCLC) (Levantini et al., 2022), glioblastoma (Yuan et al., 2025), and head and neck squamous cell carcinoma (Carosi et al., 2026). In NSCLC, activating mutations such as exon 19 deletions and L858R are common therapeutic targets of EGFR tyrosine kinase inhibitors (TKIs), including gefitinib, erlotinib, and osimertinib. Due to its central role in tumor growth and therapeutic response, EGFR remains one of the most important targets in precision oncology (Tian et al., 2025).

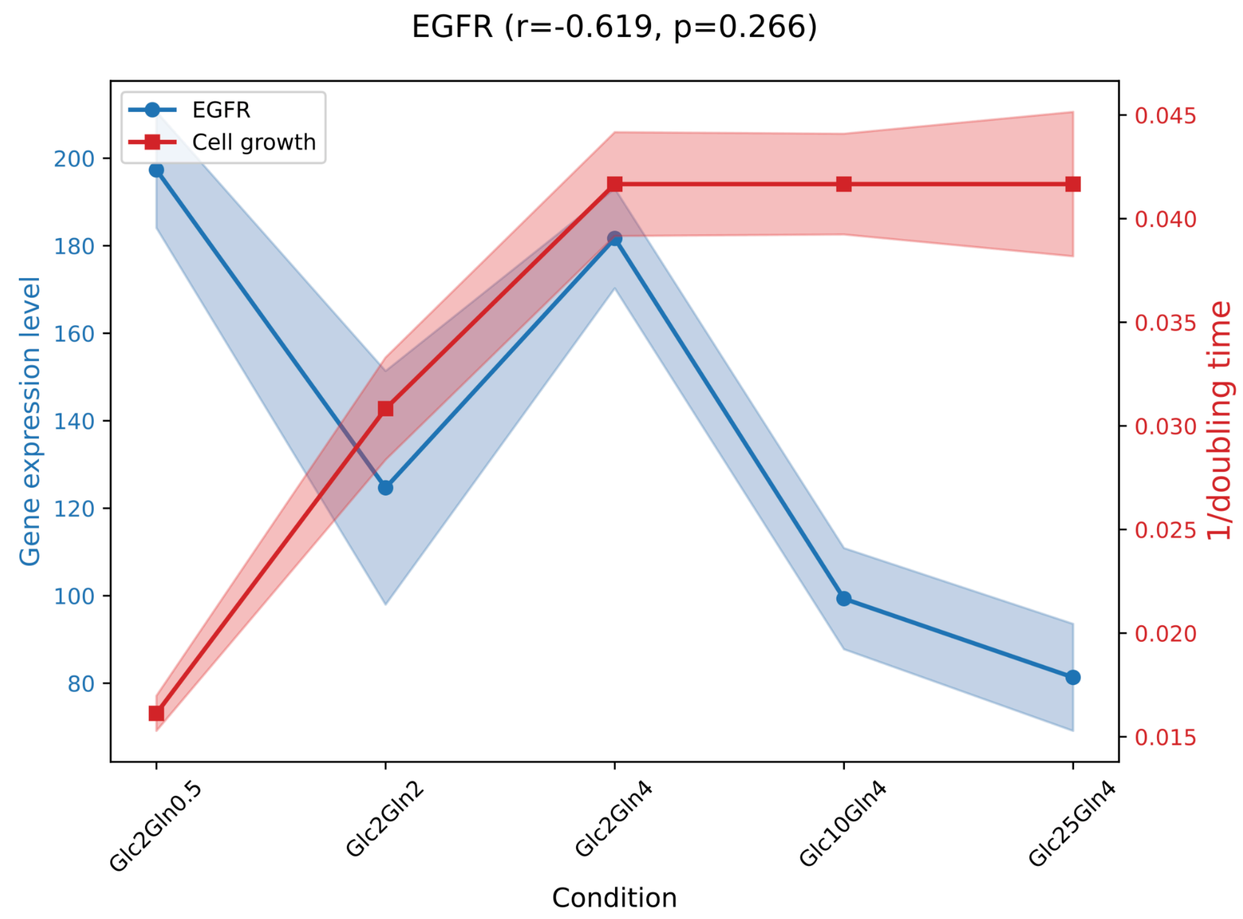


**Figure S6. The gene expression for *EGFR* under various conditions.**

### MYCL

The MYCL gene belongs to the MYC oncogene family, next to c-Myc and MYCN (Akifuji et al., 2021). It is located on chromosome 1, and its protein product weighs about 40 kDa. The family of these genes are leucine zipper transcription factors, which form heterodimers with MAX and bind to E-box elements. The basic functions include gene repression, influence on elongation during transcription, control global chromatin organization, and promote ribosomal biogenesis and protein synthesis. They also enhance the activity of Pol-I, II and III simultaneously. Compared to c-Myc, l-Myc has a much weaker effect on cell growth, apoptosis and transformation. Importantly, however, it makes the reprogramming of fibroblasts into the induced pluripotent stem cells. Moreover, MYCL expression is essential for dendritic cells, which are responsible for optimal T-cell priming. MYCL1 is amplified and overexpressed in some malignancies (Jing & Na, 2021). In small cell lung cancer (SCLC), which is a devastating neuroendocrine carcinoma, MYCL amplifications are most frequent among the MYC gene family. Studies on genetically engineered mouse models with MYCL deletion have shown the arrest of this tumor. This shows that it may be a potential target in SCLC therapy (Masso-Valles et al., 2020).

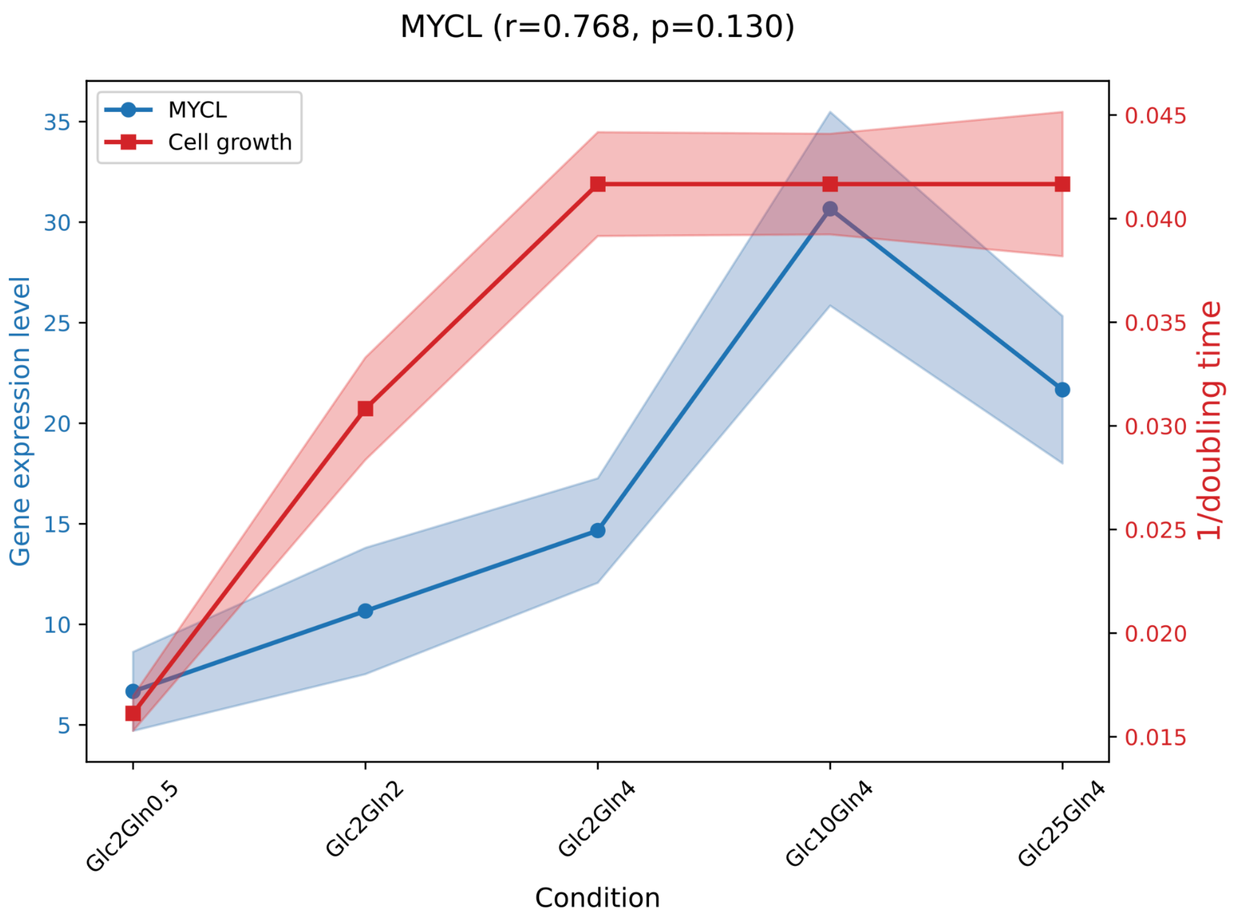


**Figure S7 The gene expression for *MYCL* under various conditions.**
